# Endophilin mediated endocytosis and Epidermal growth factor receptor govern Japanese encephalitis virus entry and infection in neuronal cells

**DOI:** 10.1101/2025.05.08.652984

**Authors:** Puja Sharma, Mukesh Tanwar, Eira Choudhary, Nandhini M. Sundaram, Divya Ojha, Laxmi Mishra, Minu Nain, Ambadas Rode, Vidya Mangala Prasad, Sudhanshu Vrati, Manjula Kalia

**Affiliations:** Regional Centre for Biotechnology, NCR Biotech Science Cluster, Faridabad, 121001, India; Molecular Biophysics Unit, Indian Institute of Science, Bengaluru, Karnataka, India; Center for Infectious Diseases Research, Indian Institute of Science, Bengaluru, Karnataka, India

**Author notes:** ICMR-National Institute of Malaria Research, Dwarka, Delhi.

**Keywords:** actin, dynamin, Fast-endophilin mediated endocytosis, flavivirus, membrane trafficking, receptor tyrosine kinase

## Abstract

Several trafficking pathways are operational at the plasma membrane, and both clathrin-dependent, and clathrin-independent endocytosis (CIE), can serve as virus entry portals. Our research has shown that the neurotropic *flavivirus*: Japanese encephalitis virus (JEV), infects neuronal cells via CIE. Here we have identified and characterized two essential host-factors for JEV trafficking in neuronal cells: Endophilin & Epidermal Growth Factor Receptor (EGFR). Through quantitative estimation of viral RNA copy number, we demonstrate that JEV entry in neuronal cells, was blocked by knock-down of Endophilin A isoforms, while activation of Endophilin-mediated endocytosis using a specific inhibitor of GSK-3β enhanced virus entry. Deletion mutants of Endophilin showed an essential role of SH_3_, BAR, H_0_ domains for virus entry. High resolution fluorescence imaging of virions showed overlap with Endophilin A2 puncta. Virus entry led to rearrangements of the actin cytoskeleton, and was highly sensitive to any pharmacological actin perturbation. Virus endocytosis activated EGFR, and the specific kinase inhibitors Erlotinib, and Gefitinib, reduced virus entry and replication in cultured cells and mouse primary cortical neurons. Silencing of EGFR, competitive inhibition with receptor ligand EGF, and EGFR specific antibodies significantly impaired JEV binding and entry, indicating the crucial role of its ligand binding domain for virus attachment/receptor interaction. EGFR colocalized with virions at early time-points of infection, and the ED3 domain of the JEV-envelope protein showed specific interaction with EGFR through BLI. Our study provides evidence for JEV entry in neuronal cells through an endocytic pathway involving Endophilin and EGFR.

## Introduction

Virus binding and entry into the host cell are the two key early determinants of infection. This involves virus interaction with attachment factors and specific receptors/co-receptors, followed by either fusion at the plasma membrane or endocytosis. Viruses can exploit multiple endocytic pathways and also activate signaling pathways that favour infection (1,2). Clathrin mediated endocytosis (CME), one of the best studied and characterized pathways in diverse physiological and disease contexts, has also been described as an entry portal for several viruses (3,4). Over the past two decades, several studies have advanced our understanding of clathrin independent endocytosis (CIE) both in terms of the molecular players and cargoes (5–7). Not surprisingly, viruses have also been shown to hijack these CIE pathways.

The focus of our study is the mosquito borne *flavivirus*: Japanese encephalitis virus (JEV), which is the leading cause of virus-induced encephalitis in South and Southeast Asia, leading to approximately 50,000 to 175,000 cases per year. The virus is neurotropic with high morbidity and mortality (8,9). JEV can infect diverse cell types and mammalian hosts, and several proteins such as Dendritic cell-specific intercellular adhesion molecule-3-grabbing non-integrin (DC-SIGN), Glucose-regulated protein 78 (GRP78), T-cell immunoglobulin and mucin domain 1 (TIM-1), heat shock protein 70, integrins and vimentin have been described as receptor molecules (10–15). Virus binding is likely to be multivalent, and other potential receptors might also be involved. The entry pathway for JEV is established to be cell-type dependent. Studies from our and other groups have shown that JEV entry in neuronal cells follows CIE (16–19), as opposed to CME in cell types such as fibroblasts and epithelial cells (20–22). An siRNA-based membrane trafficking screening study in the human neuronal cell line IMR-32 showed that JEV infection relied majorly on proteins of the actin cytoskeleton: the ARP2/3 complex, N-WASF family proteins and Rho-GTPases, and on those of the Ephrin-mediated signaling/receptor protein tyrosine kinase (RTK) signaling pathway (17). Another membrane trafficking screening study in human neuronal cell line SK-N-SH showed an important role of caveolin-1, and the EGFR-PI3K signaling pathway, leading to the activation of RhoA-mediated actin polymerization to be essential for JEV infection (19).

The cell biology of JEV internalization through CIE in neuronal cells is still not completely understood. Recently, a class of BAR domain (Bin, Amphiphysin, Rvs) family proteins have emerged to be essential for CIE pathways. One of the BAR domain family members, endophilin A, was shown to control an endocytic pathway independent of clathrin, named Fast endophilin-mediated endocytosis or FEME (23). FEME activation is triggered by ligand binding and leads to CIE of several growth factor receptors, such as the G-protein-coupled receptor (GPCR) superfamily and the RTKs (23,24). Clathrin independent and endophilin mediated endocytosis have been described in neuronal cells including synapses. Major molecular players of FEME, such as dynamin, cholesterol, actin, Rho, Rac and PAK1, are also involved in JEV entry in neuronal cells (16,17). JEV is also known to be associated with filopodia, an active site for FEME (16).

Studies have shown that multiple viruses co-opt for activation of PI3K and MAPK signaling pathways downstream of RTKs to gain entry and modulate host defense response (25). One receptor belonging to this family is the Epidermal growth factor receptor (EGFR), which is well studied in terms of the virus life cycle as an essential host factor. Multiple viruses, such as Influenza A virus, Hepatitis E virus, Hepatitis B virus, and Hepatitis C virus utilize EGFR as an entry co-factor to mediate virus internalization inside the host cell (25,26). Other viruses exploit this kinase’s complex dynamic signaling network by activating downstream signaling cascades such as EGFR/ERK/AKT to gain entry. JEV is known to exploit EGFR-PI3K signaling to activate the F-actin polymerization leading to JEV internalization (19).

This study further delineates the pathway exploited by JEV to gain entry into its target neuronal cells. We observe that endophilin A is a major host factor for JEV entry in human neuronal SH-SY5Y cells. Genetic knockdown of endophilin A isoforms leads to a decrease in virus internalization, while specific activation of the FEME pathway using an inhibitor of GSK3β enhanced virus entry. Using dominant negative domain mutants of endophilin, the crucial role of membrane curvature sensor domain BAR, and the receptor binding domain SH_3_ in JEV entry was established. JEV internalization also led to extensive remodelling of the actin cytoskeleton. Endophilin-mediated endocytosis has also been reported to be crucial for the uptake of many RTKs, such as EGFR. JEV internalization led to the phosphorylation of EGFR, and inhibition of EGFR kinase activity blocked virus entry. Treatment of cells with EGFR extracellular domain binding antibody, cetuximab, and EGF led to a decrease in JEV attachment and internalization, further supporting the hypothesis that EGFR could potentially be involved in the initial binding of JEV with the host cell surface. A specific interaction of the JEV-envelope domain (ED) III with EGFR was confirmed through BLI. Collectively, our study demonstrates the cellular trafficking and signaling pathways exploited by JEV, further deepening the understanding of the molecular pathogenesis of JEV neuronal cell entry, which ultimately will be crucial for developing suitable antiviral therapies.

## Results

### JEV binding, entry and infection in neuronal cells is clathrin-independent

JEV endocytosis was analyzed through quantitative estimation of viral RNA copy number at early times of infection using two different assay conditions (Fig 1a). One was a widely used ice-synchronized technique that involves virus binding to cells on ice for 1h, followed by washing and further incubation in complete serum medium at 37°C for 1h. However, since ice incubation has been reported to reduce membrane fluidity, and can potentially inhibit fast endocytic processes (23), we also employed an alternate assay where virus binding and entry was performed at 37°C (Fig 1a). Quantitative estimation of viral RNA was performed to analyze virus binding and endocytosis under both conditions.

**Fig 1:**
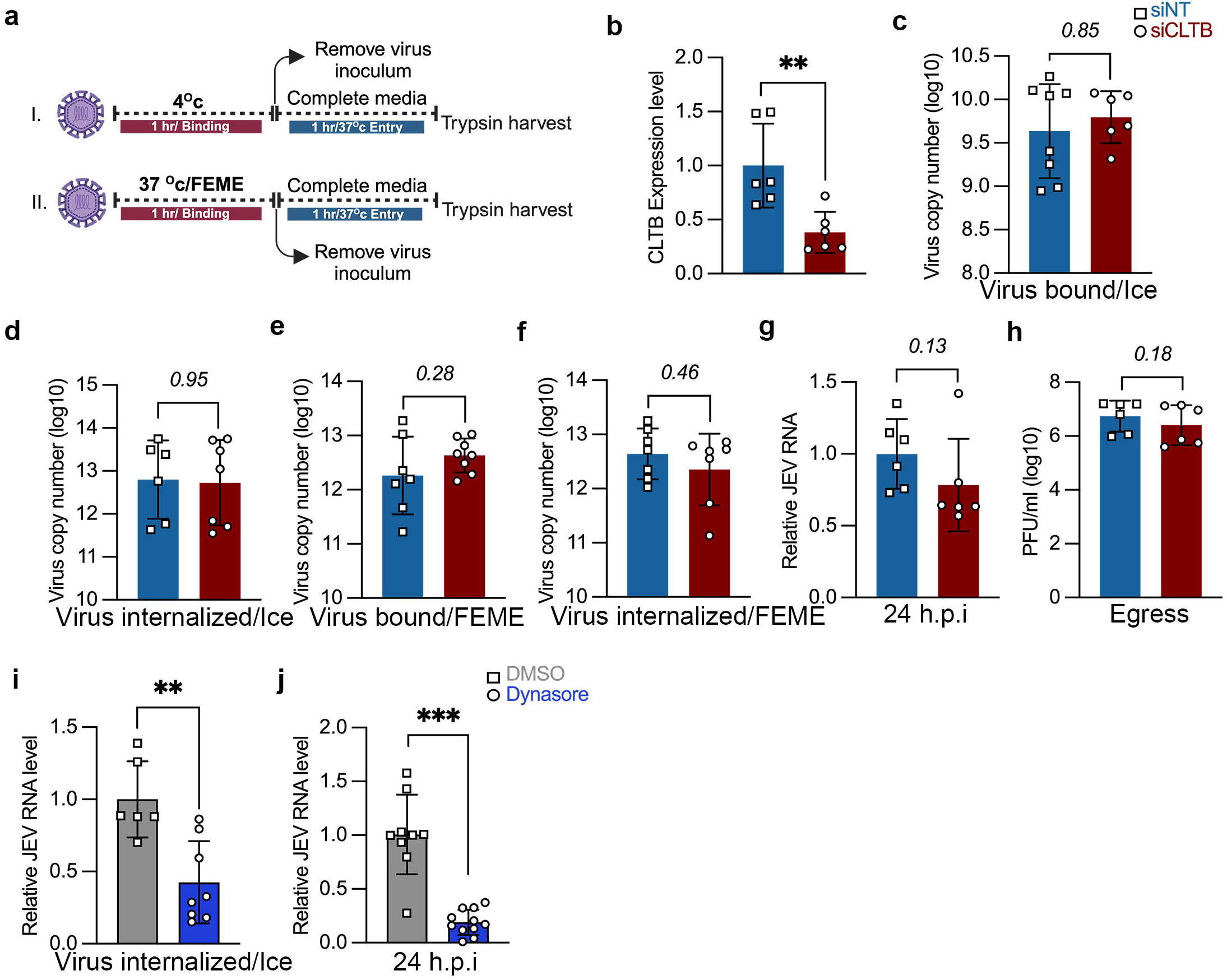
JEV binding, entry and replication in neuronal cells is clathrin-independent. (a) Schematic representation of virus binding/entry assay. (b-h) SH-SY5Y cells were transfected with siNT/si CLTB for 72 h. (b) Bar graph shows the relative level of CLC mRNA post-siRNA transfection. (c-f) The siRNA-transfected cells were infected with JEV (5/10 MOI), and virus binding/entry assays were performed as described. Absolute virus envelope copies were detected with qRT-PCR. (g-h) The siRNA-transfected cells were infected with JEV (1 MOI) for 24 h. Intracellular viral RNA levels were determined by qRT-PCR (g), and the extracellular virus particles were detected with plaque assays (h). (i) Cells seeded in nu-serum containing media were pre-treated with either DMSO or 80µM dynasore for 1 h 37°c and were infected with purified JEV as described previously. Internalized virus load was estimated by qRT-PCR, when incubated on ice (i), viral RNA levels at 24 hpi (j). All values are represented from two or more independent experiments with mean ± S.D. Statistical analysis was determined with Mann Whitney test with a 95% confidence level, NEJM: 0.12 (ns),0.033 (*), 0.002(**), <0.001(***).

To examine the role of CME in JEV entry in the human neuroblastoma cell line SH-SY5Y, an RNA interference-based knockdown approach was utilized. The effect of CLC knockdown was first tested on the internalization CME cargo: Tf and EGF. While Tf endocytosis is strictly through CME, EGF follows different endocytic routes, with a lower dose resulting in uptake through CME, whereas, at a higher dose, a major fraction of EGF gets internalized through CIE (27,28). Treatment of SH-SY5Y cells with CLC siRNA resulted in a significant decrease in the transcript levels compared to siNT, indicating efficient knockdown (Fig 1b). These cells were given a pulse of Tf (20 ng/ml) and different concentrations of EGF (10, 20, 50, & 180 ng/ml) for 5 min at 37°c. As expected under conditions of CME inhibition, in comparison to siNT, the siCLTB transfected cells showed a significant decrease in the uptake of Tf (Fig S1a-b). The knockdown of CLTB also decreased the internalization of EGF at a concentration range between 10-50 ng/ml, suggesting a role of CME in EGF uptake at these concentrations. However, a higher dose of 180 ng/ml EGF showed no perturbation in uptake, confirming its endocytosis via CIE (Fig S1a-b).

After the cargo uptake inhibition was established upon CLTB knockdown conditions, we next checked its effect on JEV endocytosis. In one set of experiments, the virus (5 MOI) was allowed to bind on the cell surface at 4°c (ice) for 1 h, which synchronizes virus binding on cells but prevents internalization. After 1 h of binding, virus entry was initiated by adding complete media and shifting the temperature to 37°c for 1 h. Cells are then harvested following trypsin treatment, which removes the extracellular bound virus particles to enable quantification of internalized viral load only (Fig 1a, upper panel). Virus binding (ice) remained unaltered, indicating that virus attachment is unaffected by CLC depletion (Fig 1c). Virus endocytosis also remained unaltered (Fig 1d).

In a parallel set of experiments, virus (10 MOI) binding was performed at 37°c for 1 h, followed by further incubation in complete medium for 1 h to permit virus entry (Fig 1a, lower panel). These physiological or FEME conditions were also used since ice incubation is known to change membrane fluidity, which can inhibit certain CIE pathways (24). The virus binding at physiological/FEME conditions showed no significant change (Fig 1e), and further virus internalization under these conditions was also unperturbed (Fig 1f). The depletion of CLTB also had no effect on virus replication as no change in the viral RNA levels was observed at 24 h (Fig 1g), along with unaltered virus titers (Fig 1h). These experiments indicate that JEV internalization in neuronal cells can follow a clathrin-independent pathway.

To test the requirement of dynamin for the endocytosis of JEV in SH-SY5Y cells, Dynasore, a cell-permeable and reversible GTPase inhibitor that inhibits the activity of dynamin was used (29). Cells were treated with 80 µM of dynasore, and given a pulse of Tf (20 ng/ml) and EGF (50 ng/ml) for 5 min at 37°c. As expected, dynasore inhibited both EGF and Tf uptake (Fig S1c). To confirm the role of dynamin in JEV endocytosis, virus entry assays was performed in the presence of the dynasore. This resulted in a ∼60 % decrease in virus internalization (Fig 1i), and a significant inhibition of virus replication (∼80 % reduction in viral RNA levels, at 24 hpi) compared to DMSO treated cells (Fig 1j). In agreement with earlier published work from our lab (16), these results demonstrate that JEV entry is dynamin-dependent.

### Endophilin is an essential host-factor for JEV entry and replication

To characterize other molecular players for JEV endocytosis, we tested the role of endophilin. Endophilin A exists in three isoforms: A1/A2/A3. Firstly, we checked the basal protein level expression of all three endophilin A proteins in three different cell types: (1) human neuronal cell line SH-SY5Y, (2) HeLa cell line and (3) mouse brain tissue. As shown earlier (24,30), endophilin A2 expression was ubiquitously observed in all three samples (Fig 2a, upper panel). In contrast, endophilin A1 expression was only observed in brain tissue, while there was lower expression of endophilin A3 in SH-SY5Y, compared to HeLa cells and mouse brain (Fig 2a, upper panel). We then utilized a genetic knockdown-based approach to establish the effect of endophilin A depletion on the uptake of Tf and EGF. We performed a triple knockdown (TKD) of all three endophilin isoforms, as in the absence of one, the other endophilin A isoforms have been shown to compensate (23). Western blots confirmed efficient knockdown by siTKD transfection (Fig 2a, lower panel).

**Fig 2:**
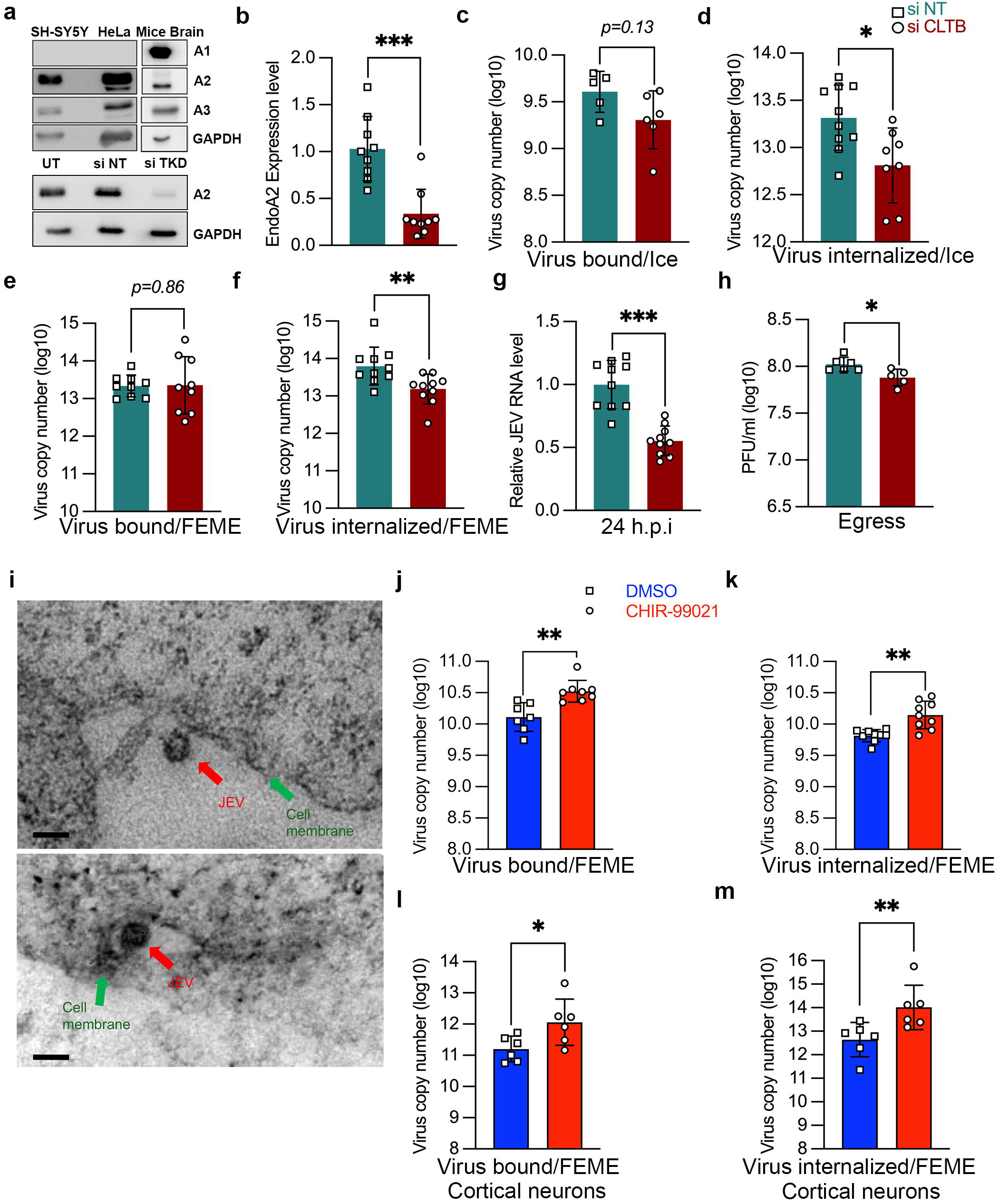
Endophilin is an essential host-factor for JEV entry and replication. (a) Protein lysates from SH-SY5Y, HeLa and homogenized mouse brain tissue were immunoblotted with endophilin A1/A2/A3 and GAPDH (internal control), upper panel. Lower panel shows levels of endophilin A2 in untransfected, NT siRNA and siTKD post-transfected SH-SY5Y cells. (b-h) SH-SY5Y cells were transfected with all three endophilin A1/A2/A3 siRNA (siTKD) for 72 h. (b) Bar graph shows the relative endophilin A2 levels in siNT/siTKD transfected cells. (c-d) Endophilin siTKD cells were allowed to bind with 10 MOI of virus on ice, and virus binding/internalization assays were performed as described previously. Absolute virus envelope copies were detected with qRT-PCR. (e-f) The siNT and siTKD transfected cells were infected with 10 MOI of the virus at 37°c for 1 h, and virus binding/internalization assays were performed. Absolute virus envelope copies were detected with qRT-PCR. (g-h) Endophilin siTKD cells were infected with 1 MOI of the virus at 37°c for 1h and were washed and harvested at 24 hpi, viral RNA was determined by qRT-PCR (g), and the extracellular virus particles were detected with plaque assays (h). (i) Representative room temperature ultramicrotomy cell sections imaged under TEM showing JEV virion localization near deeper inundations on cell surface; scale: 100 nm. (j-m) SH-SY5Y cells (j-k), or primary cortical neurons (l-m), were pre-treated with CHIR-99021 for 1 h at 37°c, and then were infected with 10 MOI of JEV at 37°c for 1h, and virus binding/internalization assays were performed as described. Absolute virus envelope copies were detected with qRT-PCR. Data shown are from two or more independent experiments represented as mean ± S.D. Statistical analysis was determined with Mann Whitney test with 95% confidence level, NEJM: 0.12 (ns), 0.033 (*), 0.002(**), <0.001(***).

SH-SY5Y cells transfected with siNT or siTKD were subjected to a pulse of EGF and Tf for 5 min at 37°c. Depletion of endophilin led to no substantial reduction in the uptake of Tf (Fig S2a-b). An earlier study (23) confirmed that endophilin A is not required for Tf uptake through CME. A dose-dependent pulse of the siTKD transfected SH-SY5Y cells with EGF (50 & 180 ng/ml) for 5 min at 37°c showed a reduction in EGF uptake at a higher concentration of 180 ng/ml but had no effect at a concentration of 50 ng/ml. These results indicated that while endophilin TKD did not affect CME, it inhibited the CIE of EGF at higher doses (Fig S2a-b). The CIE of EGF has also been shown in several independent studies in multiple cell lines such as BSC1, RPE1 and HeLa (23,31–33).

The impact of genetic inactivation of endophilin A (siTKD) was subsequently checked on virus endocytosis with entry experiments performed at two different temperatures, as stated previously (Fig 1a). The relative mRNA expression of endophilin A2 was checked in siTKD transfected cells in every experiment to confirm the knockdown at 72 h (Fig 2b). Depletion of endophilin did not affect virus binding on ice (Fig 2c). Still, it significantly reduced virus entry, validating endophilin’s crucial role in virus internalization (Fig 2d). We also performed the binding/internalization assays at a physiological temperature of 37°c. Again, in the TKD cells, the virus binding at 37°c remained unaltered (Fig 2e), while virus internalization showed significant inhibition (Fig 2f). These results were also validated to check whether this observed decrease in virus entry would manifest in lower virus replication. Indeed, we observed a substantial decline (∼55%) in viral RNA levels (Fig 2g) and virus titers (Fig 2h) at 24 hpi. These results highlight the crucial role of endophilin in JEV entry and infection.

We next checked for the colocalization of endophilin A2 with virus particles stained with specific capsid antibody, that will be indicative of internalized virions at early times of infection (before initiation of virus replication). While no signal was detected in mock-infected cells, several distinct puncta were visualized in cells where virus was added (Fig S3a). A quantitative estimation of virus particles showed that ∼70 puncta were detected per cell at 10 mpi, a time-point where virus endocytosis is initiated (Fig S3a, b). We observed colocalization of endophilin A2 with JEV particles (capsid structures) at 10 mpi (Fig S3a). Spot-spot colocalization analysis of the confocal micrographs showed ∼40 % of JEV was colocalized with endophilin A2 positive structures (Fig S3b). These data suggest that the earliest endocytic structures containing internalized JEV particles are also positive for endophilin. We also performed TEM imaging of sectioned SH-SY5Y cells to examine the early entry events of virus infection. Spherical virus particles of ∼ 50 nm were observed on the surface of SH-SY5Y cells, showing interactions in large inundations of the cell membrane that appear distinct from the classical clathrin-coated vesicles (Fig 2i).

### JEV entry is enhanced by GSK3β inhibition

To further investigate if there are any parallels between endophilin-mediated CIE of JEV and the FEME pathway, we treated SH-SY5Y cells with a small molecule compound CHIR-99021, a well-reported kinase inhibitor of GSK3β, which suppresses FEME under normal physiological conditions (34). Treatment of cells with CHIR-99021 for 1 h at 37°c enhanced the uptake of EGF, further supporting the observation that inhibition of GSK3β leads to upregulation of FEME (Fig S4a). The effect of this inhibitor was then checked on virus binding/internalization at a physiological temperature of 37°c (FEME conditions). Treating cells with CHIR-99021 led to a significant increase in virus binding and internalization (Fig 2j-k). These results suggest that relieving the autoinhibition of FEME by treating cells with inhibitors of kinases that regulate the key steps of the pathway substantially increased JEV endocytosis. These findings were further corroborated in the primary culture of cortical neurons harvested from mice brain, which showed a similar enhancement of JEV entry on the CHIR-99021 treatment (Fig 2l-m).

Next, we tested whether JEV endocytosis increases the fluorescence intensity of endophilin-positive structures that are a hallmark of FEME (23,24). Since serum stimulation has been shown to increase FEME (34), we used it as a positive control along with EGF stimulation. Primary cortical neurons were stimulated with 20% serum for 15 min at 37°c. Compared to the control, serum stimulation increased endophilin-positive structures, which can be observed as discrete punctate structures when immunostained with endophilin A2. An EGF pulse (180 ng/ml) for 5 min at 37°c also increased endophilin-positive structures. JEV infection also enhanced the endophilin-positive punctate structures at 15 mpi, which further increased at 30 mpi. (Fig S4b-c). This suggests that like serum stimulation, high EGF doses, and JEV endocytosis can increase endophilin-positive structures, likely indicative of an upregulation of the FEME pathway. Collectively, our data on the genetic depletion of endophilin and molecular upregulation of FEME suggest a prominent role of this pathway in JEV endocytosis.

### Endophilin domain mutants block JEV internalization and replication

To further characterize the role of endophilin A2 for JEV internalization, we generated stable clones expressing EGFP tagged endophilin A2 full length (FL), or the domain truncated mutants ΔSH_3_, ΔH_0_, and ΔBAR in SH-SY5Y cells (Fig 3a). The over-expression of the endophilin domain truncated mutants will impose a dominant effect and override the endogenous protein function. The over-expression of the constructs was confirmed through immunoblotting with anti-GFP antibody. Western blot showed bands at the expected molecular weights for FL and truncated domains of endophilin A2 (Fig 3b). Confocal microscopy images show GFP expressing stable cells with EGFPC1, FL, ΔSH_3_, ΔH_0_, and ΔBAR overexpression (Fig 3c).

**Fig 3:**
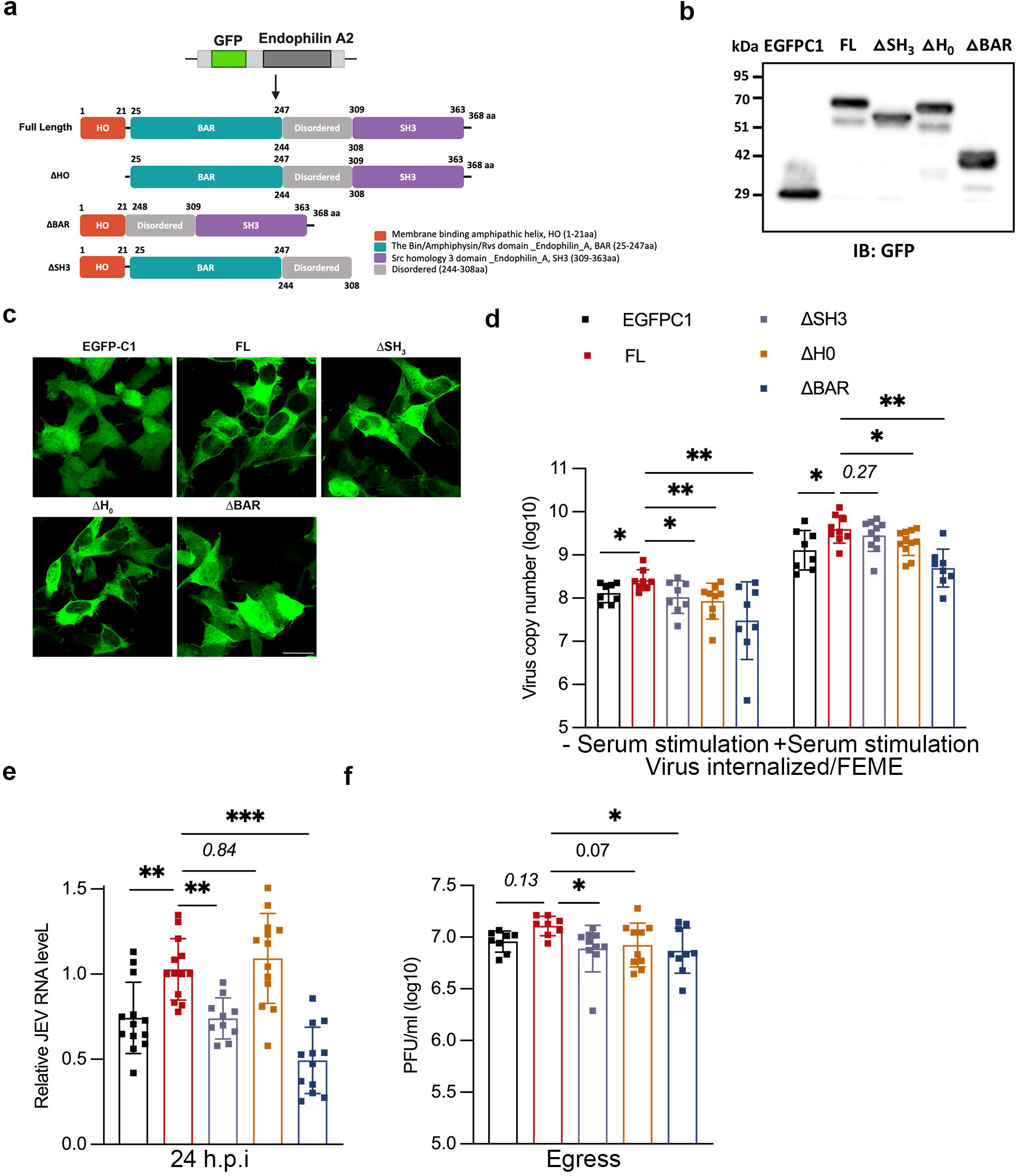
Endophilin domain mutants block JEV internalization and replication. (a) Schematic representation of the GFP endophilin A2 with full length (FL), ΔSH_3_, ΔH_0_, and ΔBAR domains deletion mutants. (b) Lysates of SH-SY5Y cells stably expressing EGFPC1, FL, ΔSH_3_, ΔH_0_, and ΔBAR domains were immunoblotted with anti-GFP antibody. (c) Confocal micrographs of cells stably expressing the various domains. Scale: 10 µm (d) Endophilin domain mutant cell lines were stimulated with or without 20% FBS for 1 h at 37°c, infected with 10 MOI for 1 h at 37°c, and harvested post 1 h with trypsin. The histogram represents the viral copy number of internalized virus particles. (e-f) Cells were infected with 1 MOI of JEV for 1 h and were harvested post-24 h. Viral RNA was determined by qRT-PCR (f), and the extracellular virus particles were detected with plaque assays (g). All values are represented as mean ± S.D from at least two independent experiments. Statistical analysis was determined using ordinary one-way ANOVA with Dunnett’s multiple comparison test with a 95% confidence level. Statistical significance: NEJM: 0.12 (ns),0.033 (*), 0.002(**), <0.001(***).

These clones were tested for cargo uptake assays to demonstrate their effect on EGF uptake at low and high concentrations. An increase in the uptake of fluorescently labelled EGF was observed in the cells expressing FL endoA2 as compared to EGFPC1 at both concentrations, indicating upregulation of the pathway upon endophilin A2 over-expression. Cells with the domain truncated mutants of ΔSH_3_, ΔH_0_, and ΔBAR displayed a decrease in the uptake of EGF (Fig S5a first & second panel, b-c). These mutants specifically perturb endophilin mediated endocytosis and not CME, as the uptake of Tf remains unchanged (Fig S5a third panel, d). The impact of these stably overexpressing cells was also checked on the fluid-phase uptake of FITC-dextran, which was unaltered (Fig S5a fourth panel, e). These data demonstrate that the endophilin A2 domain deleted mutants exert a dominant negative effect in blocking EGF uptake in SH-SY5Y cells.

We next checked for the role of each of these domains on JEV internalization. The virus entry assays were next performed under FEME conditions: 37°c for 1 h to allow virus binding at 10 MOI, with subsequent internalization at 1 h at 37°c. Here, an increase in virus entry was observed with FL overexpressing cells compared to the empty vector. A subsequent decline in the virus entry was observed with all three truncated mutants, with a maximum decrease with ΔBAR-expressing cells. This result highlights the crucial role of the three main domains of endophilin A2 proteins in JEV entry (Fig 3d, left panel). To further strengthen our results, we checked virus entry in these mutant cell lines when stimulated with additional serum (20% FBS) to enhance FEME. Entry experiments showed an increase in virus copy numbers with FL compared to the empty vector, while the ΔH_0_ and ΔBAR mutants showed a decrease (Fig 3d, right panel). We also checked the effect of these endophilin mutant cell lines on virus replication. As previously observed with our entry experiments, we observed an increased viral RNA level in cells over-expressing FL compared to empty vector. The ΔSH_3_ and ΔBAR-expressing cells showed a reduction in virus replication as compared with the FL, and maximum inhibition was observed with ΔBAR deletion (∼50%) (Fig 3e). These results were also supported by reduced virus titers in ΔSH_3_ and ΔBAR domain-expressing cells (Fig 3f). Collectively our data shows that over-expression of endophilin A2 enhances JEV uptake and also establishes a crucial role of the cargo receptor binding domain and the membrane curvature-inducing domains of endophilin A2 for JEV internalization.

### Pharmacological inhibition of actin cytoskeleton perturbs JEV entry and replication

Actin polymerization is crucial for multiple CIE pathways. The molecular mechanism of FEME is known to be an actin-driven pathway, vital for carrier formation and driving the vesicles into the cytoplasm. We checked for the role of the actin machinery in JEV entry through pharmacological inhibition of the pathway using various known inhibitors, Latrunculin A (Lat A) which inhibits actin polymerization (35), Jasplakinolide (Jas) which binds to F-actin and blocks the actin depolymerization (36), and Cytochalasin D (CytoD) which induces the depolymerization of actin filaments through binding to the G-actin leading to dimer formation and nucleation of the newly formed filaments (37,38). The other small molecule inhibitors used were CK-548, which binds to the Arp2/3 complex and inhibits its ability to polymerize actin, and Wiskostatin, an N-WASP-mediated actin polymerization inhibitor (39). Cell viability assays were performed to establish non-toxic drug concentrations: LatA (1uM), Jas (1uM), CytoD (5uM), CK-548 (50uM) and Wiskostatin (10uM) (Fig S6a). Cells pre-treated with the drugs for 1 h inhibited virus binding and internalization at 37°c (Fig 4a, b). This was also observed through confocal imaging, wherein treatment of cells with the drugs led to a decrease in virus endocytosis, which can be observed with a reduced number of envelope positive immunostained structures (Fig 4c, d). Treatment of cells with CytoD, CK-548 and Wiskostatin inhibited virus replication as observed by a decrease in viral RNA levels (Fig S6b) and virus titers (Fig S6c), thus confirming the critical role of a functional actin machinery for JEV infection.

**Fig 4:**
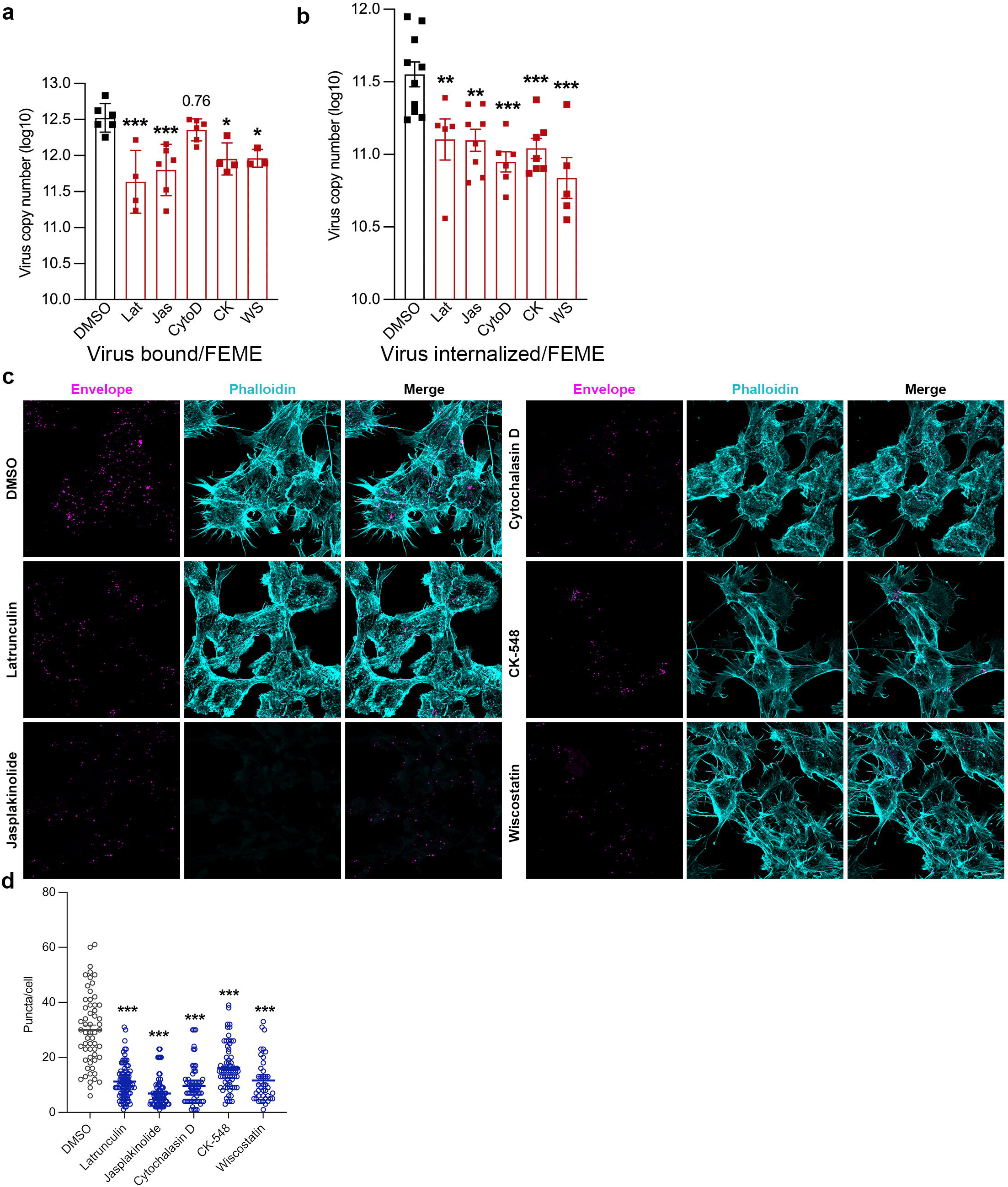
Pharmacological inhibition of actin cytoskeleton perturbs JEV entry and replication. (a-b) SH-SY5Y cells were pre-treated with DMSO control, LatA (1uM), Jas (1uM), CytoD (5uM), CK-548 (50uM) and Wiskostatin (10uM) for 1 h at 37°c, followed by 10 MOI JEV at 37°c for virus binding. Cells were harvested with trypsin treatment to remove extracellular virus particles. Absolute virus envelope copies were detected with qRT-PCR. (c) SH-SY5Y cells were pre-treated with DMSO control/ drug for 1 h at 37°c subsequently followed with 100 MOI virus binding at 4°c. Cells were then shifted to 37°c for virus internalization for 15 min, fixed, immunostained with Alexa Fluor 546-phalloidin (cyan), and viral envelope particles (magenta) and imaged with 63x objective. The representative confocal micrograph shows the phalloidin-stained F-actin and JEV envelope puncta in the presence of actin inhibitors, scale: 10 µm. (d) The histogram quantifies the number of envelope positive puncta per cell analyzed through Image J. All values are represented as mean ± S.D./S.E.M; statistical analysis was determined with Ordinary one-way ANOVA/Kruskal-Wallis test with Dunn’s multiple comparisons test. Statistical significance: NEJM: 0.12 (ns),0.033 (*), 0.002(**), <0.001(***).

Through confocal microscopy, we monitored changes in the actin cytoskeleton upon JEV entry at different time points, starting from 5 min to 60 min pi. Mock-infected cells showed cortical actin meshwork fibers with different well-aligned stress fibers, including dorsal and peripheral fibers (Fig S7a, uninfected panel). In the JEV-infected cells, by 5 min pi, the F-actin redistribution changed, and dense condensed F-actin filaments could be observed with long protrusions in the filopodia (Fig S7a, 5 min panel). By 10 min post-virus entry, increased and dense lamellipodia structures were observed (Fig S7a, 10 min panel). Long filopodia extensions were observed at 15 min and 30 min post-virus entry (Fig S7a, 15- and 30 min panel). The number of protrusions decreased by 60 min post-virus entry (Fig S7a, 60 min panel). We also checked actin rearrangements in primary cortical neurons infected with JEV, which showed multiple long protrusions with denser lamellipodium (Fig S7b).

### EGFR signaling is activated and essential for JEV entry

Several viruses such as ZIKV, PEDV, and IAV, can activate EGFR signaling cascade (40–42). JEV has also been shown to activate EGFR-PI3K signaling during entry (19). To test the same in our experimental setup, SH-SY5Y cells were infected with purified JEV, and the levels of phosphorylated EGFR were analyzed through western blotting at 0, 5, 10, 15, 30, and 60 min of virus internalization (Fig 5a). Compared to time-matched mock infected cells, phosphorylated EGFR was detected in JEV-infected cells peaking at 15 min pi. EGF treatment (100 ng/ml) was also used as a positive control (Fig 5a). These observations were also confirmed with confocal microscopy, wherein EGF treatment or JEV entry resulted in the activation of p-EGFR compared to mock-infected cells (Fig 5b).

**Fig 5:**
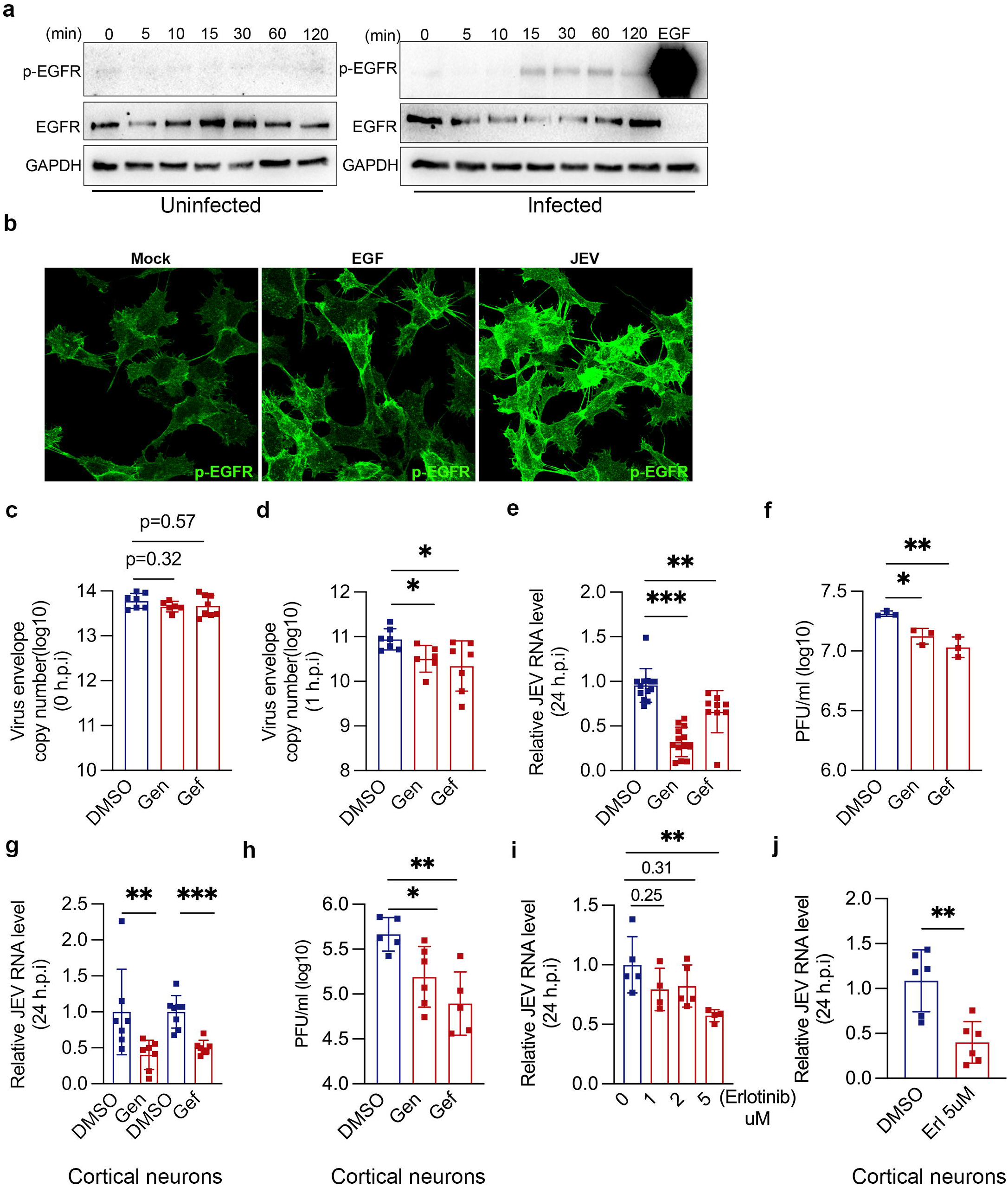
EGFR signaling is activated and essential for JEV entry. (a) SH-SY5Y cells were first allowed to bind with purified JEV at 4°c for 1h and were subsequently shifted to 37°c for 0, 5, 10, 15-, 30-, 60-, and 120-min. Cell lysates were analyzed through western blotting using p-EGFR, EGFR and GAPDH (loading control) antibodies. (b) Representative confocal images of SH-SY5Y cells: mock, treated with EGF (100 ng/ml), or JEV (100 MOI, 4°c, 1h for attachment, and then shifted to 37°c for 15 min), immunostained with p-EGFR antibody. (c-f) SH-SY5Y cells, (g-h) primary cortical neurons, were serum starved overnight and treated with genistein (400 ug/ml) or gefitinib (400 ug/ml) for 5 h before infection. (c) Virus attachment assay (10 MOI, 4°c, 1 h). (d) Virus entry assay (10 MOI, 4°c, 1 h binding and then shifted to 37°c for 1h). (e-f) Virus replication assay (1 MOI, 37°c, 24 hpi). Viral RNA was determined by qRT-PCR (c-e), and the virus titers were quantified with plaque assays (f). (g-h) JEV RNA levels and titers in primary cortical neurons (1 MOI, 37°c, 24 hpi). (i-j) SH-SY5Y cells (i), primary cortical neurons (j), were pre-treated with erlotinib for 4 h at 37°c and then infected with JEV (1 MOI, 37°c, 24 hpi). Viral RNA was determined by qRT-PCR. Data shown are from two or more independent experiments represented as mean ± S.D. Statistical analysis was determined by Ordinary one-way ANOVA with Dunnett’s multiple comparison tests or the Mann-Whitney test with 95% confidence level. Statistical significance: NEJM: 0.12 (ns),0.033 (*), 0.002(**), <0.001(***).

To examine the role of EGFR signaling in the context of JEV endocytosis, we utilized the broad-spectrum RTK inhibitor genistein, and the EGFR-specific inhibitors: gefitinib and erlotinib. To ensure that the drugs do not exert any cytotoxic effect, a cell viability assay was performed (Fig S8a, b). Serum-starved SH-SY5Y cells were pre-treated with 400 ug/ml of genistein or gefitinib for 5 h at 37°c. This did not alter virus binding or attachment (Fig 5c) but significantly reduced virus entry (Fig 5d), replication (Fig 5d), and production of infectious virions (Fig 5f). We further confirmed our results in primary cortical neurons and observed a significant reduction in JEV RNA levels and titers with genistein and gefitinib treatment (Fig 5g, h).

These observations were validated with erlotinib, another known small molecule inhibitor of EGFR tyrosine kinase activity. Erlotinib showed antiviral activity in SH-SY5Y cells with ∼50 % reduction in viral RNA levels at 24 hpi (Fig 5i). A similar inhibition in virus replication was also observed in primary cortical neurons (Fig 5j). Collectively, pharmacological inhibition of EGFR with both broad-spectrum and specific inhibitors was detrimental for virus entry and infection.

### JEV entry is impaired in EGFR depleted cells

We used siRNA targeting of the human EGFR gene to specifically knockdown its expression. SH-SY5Y cells were transfected with siNT or siEGFR, and a significant reduction in the EGFR mRNA levels was observed at 72 h of transfection compared to the control siRNA transfected cells (Fig 6a). As expected, the silencing of EGFR led to a decrease in the cargo uptake of fluorescently labelled EGF (Fig S9a-b). EGFR silencing resulted in a significant decrease in the virus binding (Fig 6b), and entry under both ice synchronization (Fig 6c), and FEME conditions (Fig 6d), suggesting the potential role of EGFR in virus attachment and entry. EGFR silencing also reduced viral RNA levels by ∼40% at 24 hpi, associated with decreased virus titers (Fig6 e-f). These results indicate the role of EGFR as an essential entry co-factor for JEV internalization in neuronal cells.

**Fig 6:**
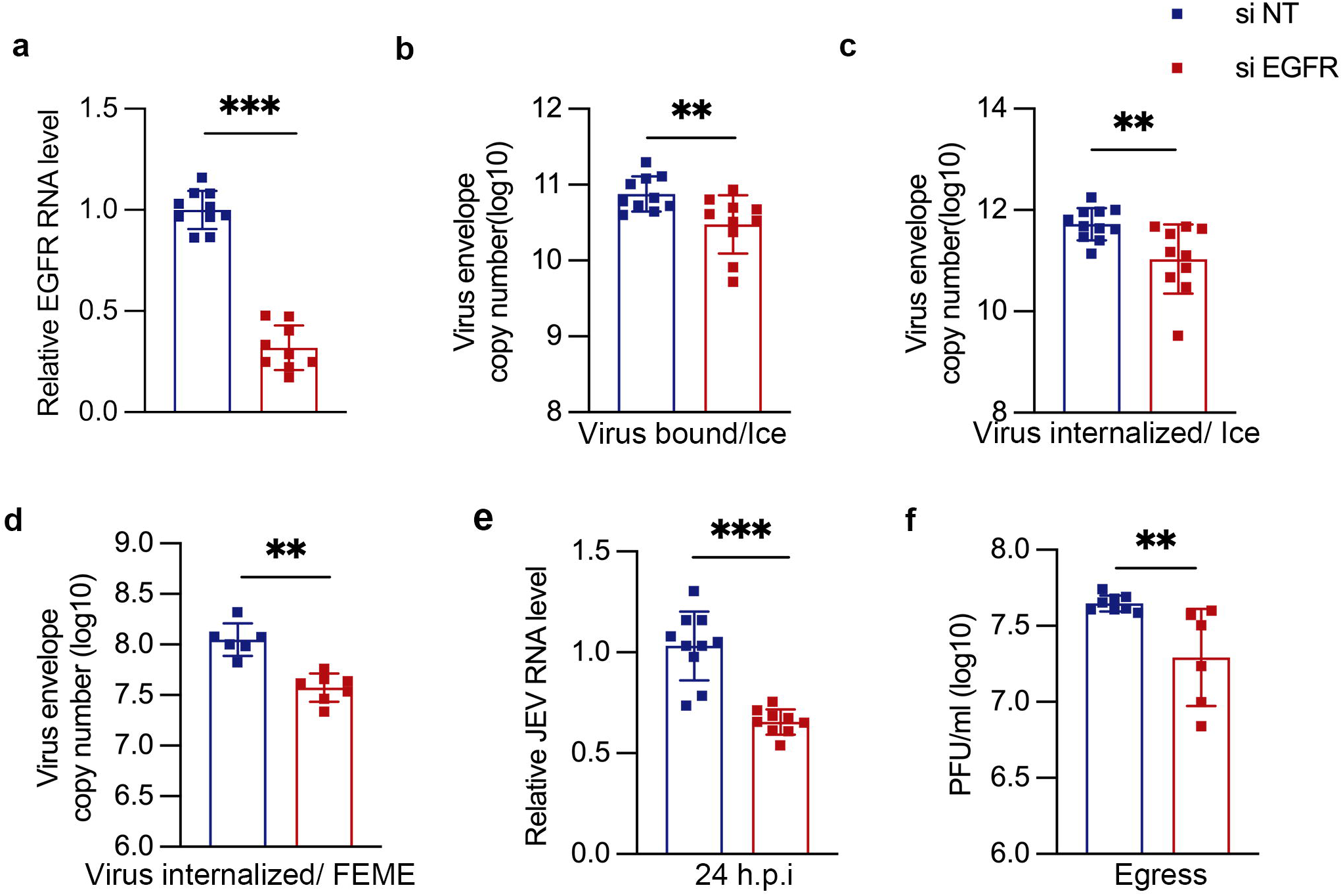
Silencing of EGFR decreases JEV entry. SH-SY5Y cells were transfected with transfected with siNT/si EGFR for 72 h. (a) Histogram shows the relative level of EGFR knockdown at the mRNA level post-siRNA transfection. (b) JEV attachment on cells (10 MOI, 4°c, 1 h). (c) JEV internalization (10 MOI, 4°c, 1h binding and internalization at 37°c for 1 h). (d) JEV internalization under FEME conditions (10 MOI, 37°c, 1 h). (e-f) JEV replication (1 MOI, 24 hpi). JEV RNA levels were determined by qRT-PCR (b-e), and titers plaque assays (f). All values are represented from two or more independent experiments with mean ± S.D. Statistical analysis was determined with Mann Whitney test with a 95% confidence level, NEJM: 0.12 (ns),0.033 (*), 0.002(**), <0.001(***).

### Ligand binding domain of EGFR plays a critical role in JEV entry

Since EGFR gene silencing inhibited virus attachment, we next explored the role of the EGFR ligand binding domain in virus entry. An antibody inhibition-based approach was used to specifically block the ligand binding domain of the receptor. Incubation of cells with an EGFR epitope binding monoclonal antibody significantly decreased virus attachment (Fig 7a) and entry (Fig 7b), as compared to isotype control. This further manifested as reduced virus replication and egress at 24 hpi (Fig 7c, d). Thus, the EGFR ligand binding domain appeared to be critical for virus attachment on cells and further infection.

**Fig 7:**
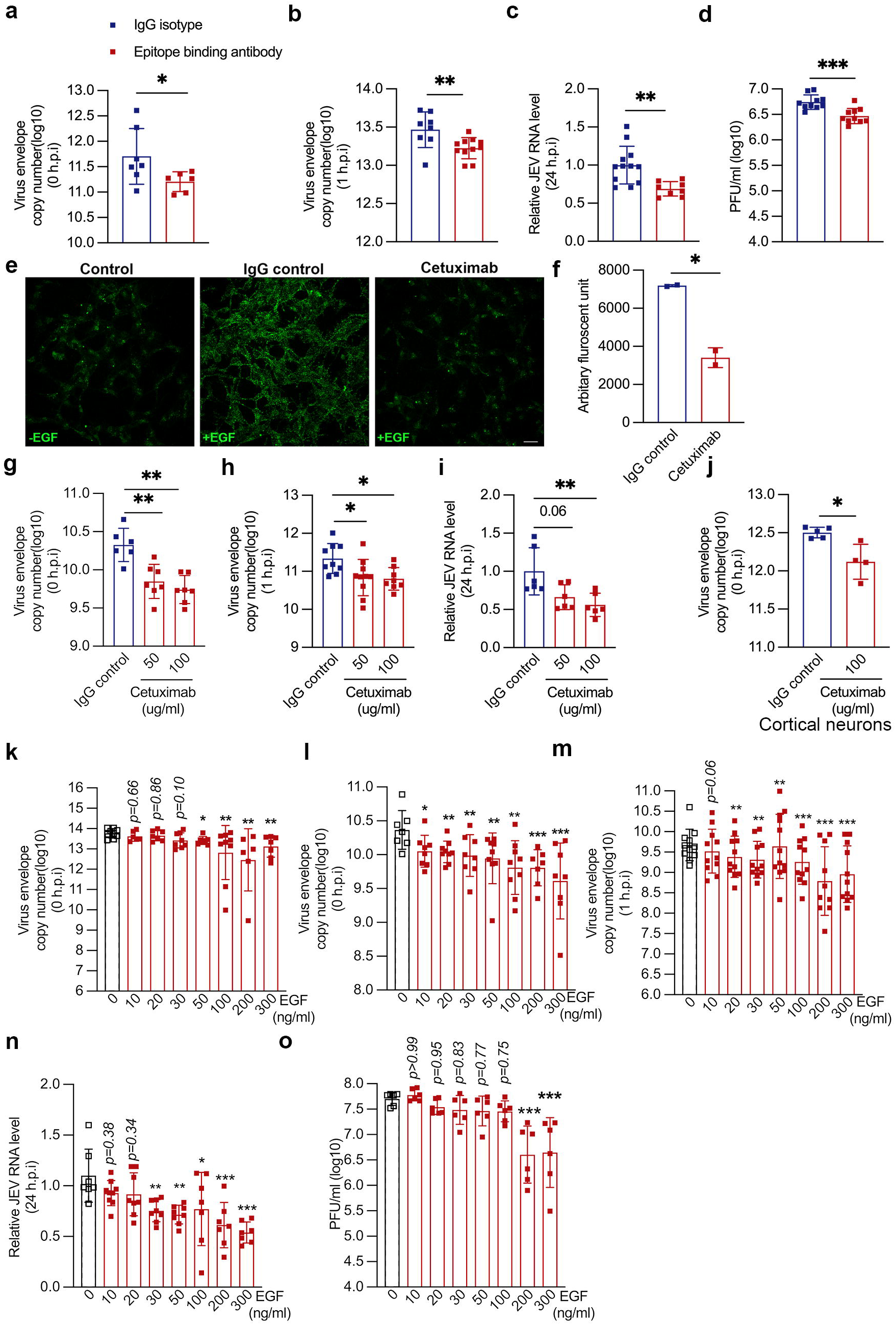
Ligand binding domain of EGFR is essential for JEV entry. (a-d) SH-SY5Y cells were pre-incubated on ice with an antibody mapping to a cell surface epitope of EGFR for 1 h. (a) JEV attachment (10 MOI on ice, 30 min). (b) JEV entry (10 MOI virus at 4°c for 30 min and then shifted to 37°c for 1 h). (c-d) JEV replication (1 MOI, 24 hpi). Viral RNA levels were determined by qRT-PCR (a-c) and titres by plaque assays (d). (e-f) SH-SY5Y cells were left untreated or incubated with IgG isotype control or cetuximab at 100 ug/ml concentration for 3 h at 37°c. Cells were then pulsed with Alexa fluor 555 EGF (100 ng/ml) for 5 mins at 37°c. (e) Representative images, scale: 20 µm. (f) Histogram quantifies the total fluorescent units of EGF uptake under untreated, IgG isotype control and cetuximab-treated conditions. Analysis was performed using image J software with ∼100 cells/coverslip, represented as mean ± S.E.M. (g-j) Cells were pre-treated with either IgG isotype control or with cetuximab (50 and 100 ng/ml) at 37°c for 3 h. (g-i) Virus attachment (10 MOI, ice, 1h) in SH-SY5Y cells (g), (h) Virus entry (10 MOI, ice 1 h, shifted to 37°c for 1 h). (i) Virus replication (1 MOI, 24 hpi), (j) Virus attachment (10 MOI, ice, 1h) in primary cortical neurons. Viral RNA levels were quantified through qRT-PCR. (k) SH-SY5Y cells were serum stimulated with 20 % FBS for 15 min at 37°c, and were allowed to bind with 10 MOI virus along with different doses of EGF at 4°c for 1 h. The histogram shows the bound viral copies detected through qRT-PCR. (l) Cells were serum-stimulated with 20 % FBS, followed by EGF stimulation at 37°c for 15 min, and then 1 MOI virus at 4°c for 1 h. (m) Serum and EGF-stimulated cells were infected with 10 MOI viruses and different EGF concentrations for 1 h on ice and were harvested post 1 h with trypsin treatment. (n) Intracellular RNA levels were detected at 24 hpi, and (o) the extracellular viral load was detected through plaque assay. All values are represented as mean ± S.D from at least two independent experiments. Statistical analysis was determined by unpaired students’ t-test or ordinary one-way ANOVA, with Dunnett’s multiple comparison test, with a 95% confidence level. Statistical significance: NEJM: 0.12 (ns),0.033 (*), 0.002(**), <0.001(***).

This was confirmed by treatment of cells with Cetuximab (Erbitux), a recombinant human chimeric monoclonal IgG1 antibody which binds to the EGF binding domain (43). Compared to the isotype control, cetuximab significantly decreased EGF uptake (Fig 7e, f). Consistent with the previous results, we also observed a reduction in JEV attachment, entry and replication in cetuximab-treated cells as compared to isotype control (Fig 7g-i). This was also validated in primary cortical neurons, which showed similar inhibition of virus binding (Fig 7j). These results indicated the possibility that JEV could be binding to EGFR at a site that overlaps with the EGF binding.

To further elucidate this, we stimulated cells with EGF and observed its effect on the early events of the virus life cycle. The binding of EGF with its receptor EGFR leads to receptor dimerization and activation of the downstream signaling cascade. However, prolonged stimulation with EGF results in ligand-induced receptor degradation (44). The effect of adding different doses of EGF was first checked on virus attachment. A stimulation step with 20 % additional serum was also performed to activate FEME. Here, we observed a decrease in virus binding starting from 50 ng/ml of EGF (Fig 7k). These results suggested that incubation of both EGF and JEV could result in a competitive inhibition for binding to EGFR. Next, we treated the serum-stimulated cells with the above-mentioned doses of EGF before infection. This step was performed to activate the receptor internalization before infection. A decrease in virus binding was observed starting from 10 ng/ml of EGF, which decreased further with an increase in the EGF dose (Fig 7l). This might result from a reduced level of surface EGFR, which is essential for virus attachment. The EGF treatment also correlated with a reduction in the virus entry at 1 hpi starting from 20 ng/ml EGF dose, which decreased further with an increase in EGF dose (Fig 7m). Reduced virus replication and titers at 24 hpi was also observed (Fig 7n, o). Altogether, our results highlight two crucial observations: first, the importance of the accessibility of EGFR ligand binding domain during the initial virus attachment, and second, the availability of the surface receptor for virus entry.

### EGFR colocalizes with virions and interacts with ED3 domain of envelope protein

Our data suggests that JEV could potentially be associating with EGFR for attachment and entry into cells. Next, we checked for interaction of EGFR proteins with JEV particles during entry. SH-SY5Y cells were allowed to bind with JEV (MOI 100) on ice, followed by internalization for 0, 10, 30, and 60 min. The virus particles can be visualized with JEV-E antibody as punctate structures (magenta) (Fig 8a). Maximum number of envelope-positive structures were observed at 0- and 5-min post entry, which then reduced at 30- and 60-min pi. The virions were observed to be colocalized with EGFR as soon as 0 min post-entry, which corresponds to the initial binding of the virus on the cell surface. The colocalization of EGFR with envelope structures then gradually reduced staring from 10-min post-entry onwards (Fig 8b). A similar overlap of JEV capsid structures with the EGFR protein was observed in SIM imaging (Fig S10a). We also checked the colocalization of the virus particles with EGFR at early time-points of virus entry in primary cortical neurons, and a similar pattern of colocalization at 0 min was observed (Fig S10b).

**Fig 8:**
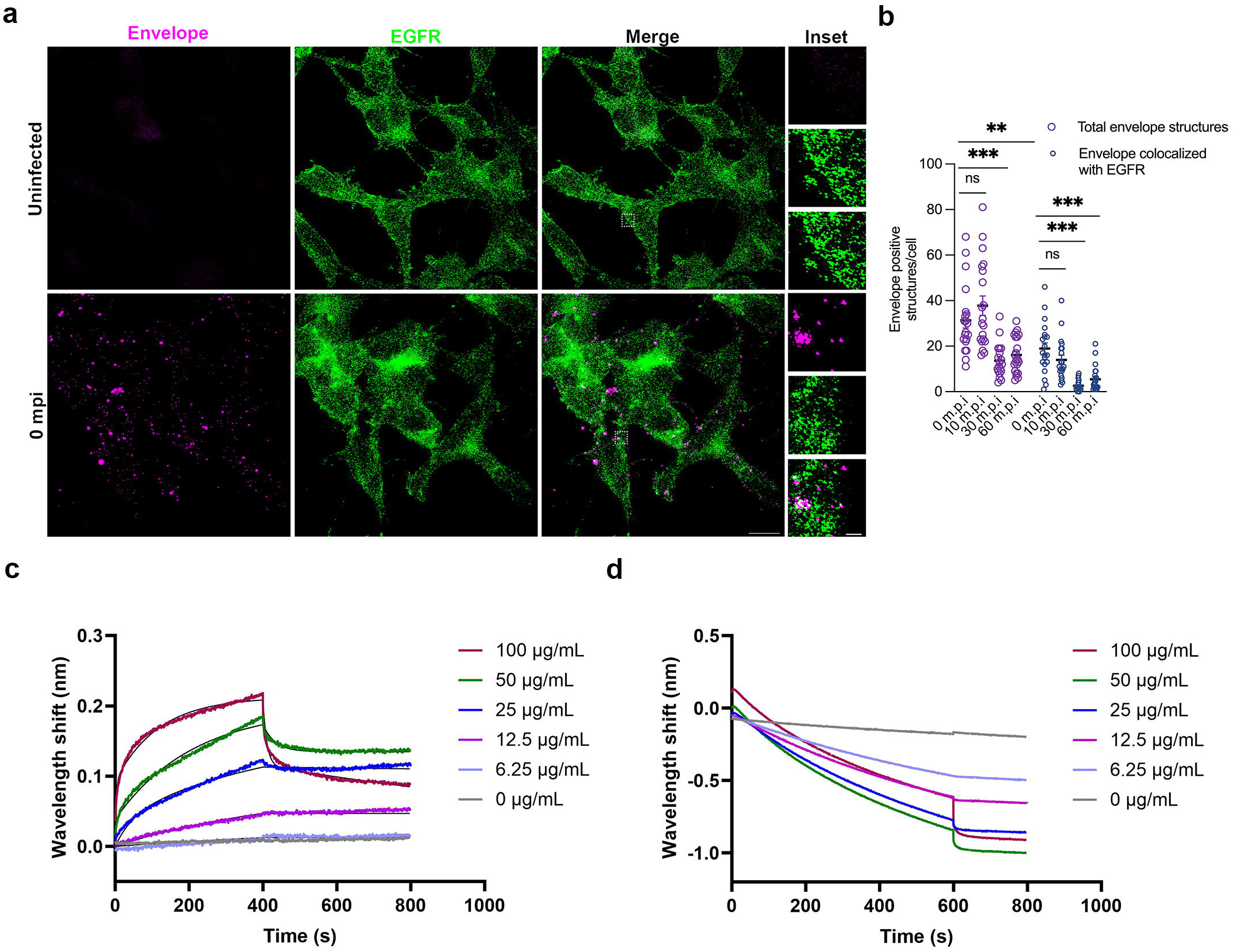
EGFR colocalizes with virions and interacts with ED3 domain of envelope protein. (a-b) SH-SY5Y cells were incubated with 100 MOI of virus for 1 h on ice and were subsequently fixed post at 0, 10, 30, and 60 mins of virus internalization. (a) Cells were immunostained with JEV envelope (magenta) and EGFR (green) antibodies. Confocal images for 0 mpi is shown for reference, other time points image not included. Insets show the zoomed image of envelope-positive structures colocalized with EGFR; scale: 10 µm, inset: 1 µm. Images were acquired in Elyra PS1 (Carl Zeiss Super-resolution microscope) using 63 x objective. (b) The histogram quantifies total envelope-positive structures and the number of envelope-positive structures colocalized with EGFR per cell at different time points. Image analysis was performed using Imaris. All values are represented as mean ± S.E.M; statistical analysis was determined with Kruskal-Wallis test with Dunn’s multiple comparisons test. Statistical significance: NEJM: 0.12 (ns),0.033 (*), 0.002(**), <0.001(***). (c-d) Bio-Layer Interferometry analysis of EGFR binding to JEV-ED3 protein. The association and dissociation curves resulting from the binding events are shown, wherein the real-time binding sensorgrams are represented in colored lines and their fitting curves are represented in black lines. (c) The JEV-ED3 protein is titrated against different concentrations of recombinant Human EGFR Protein (ECD, hFc Tag), ranging from 0-100 μg/ml. The equilibrium dissociation constants (*K_D_)* were calculated from three independent experiments using the Octet Data Analysis software’s 2:1 (Heterologous binding model) binding model. (d) The JEV-ED3 protein is titrated against different concentrations of recombinant Human EGFR Protein (Isoform VIII, hFc Tag), ranging from 0-100 μg/ml. The binding experiment was repeated three times.

The immunoglobulin-like-E protein domain III (ED3) plays a major role in receptor binding and harbors key residues that bind to neutralizing antibodies (10,45). We used the purified recombinant JEV-ED3 protein to test for specific interaction with EGFR and EGFR mutant protein vIII, which lacks the ligand binding ability with EGF or any known ligands, using Bio-layer interferometry (BLI). Based on the obtained results, it was evident that the JEV-ED3 protein shows a significant preference for EGFR compared to the EGFR mutant protein vIII (Fig. 8c, d). The *K_D_* value of JEV3 against EGFR was calculated to be 283 ± 75 nM. Collectively, these results demonstrate the fundamental importance of the EGFR extracellular ligand binding domain and the intracellular kinase domain for JEV binding and entry.

## Discussion

Viruses exploit different endocytic pathways to gain entry into the host cell. While CME remains the best characterised pathway till date, our understanding of CIE has expanded, and recent studies have highlighted the potential of these pathways as virus entry portals. This is likely to be influenced by the cell type and expression level of attachment factors and receptors. Several host proteins have been described as putative JEV receptors (15,46–49). Studies have established that JEV enters through CME into various cell types such as fibroblast, epithelial, and insect (16,20–22,50,51), while it targets a clathrin-independent pathway for entry and infection in neuronal cells (16,17,19,52).

This study characterizes the pathway exploited by JEV for entry in the SH-SY5Y human neuronal cell line, and in mouse primary cortical neurons. In line with our previous observations in mouse Neuro2a cells, we observe an active dynamin-dependent CIE pathway operational for JEV entry in human neuronal SH-SY5Y cells, that is primarily controlled by the Endophilin protein. Endophilin A has been shown to be critical regulator of a receptor-mediated clathrin-independent pathway, also described as FEME (23). Kinases such as Cdk5 and GSK3β inhibit dynein recruitment to FEME carries, and are thus negative regulators of this pathway (34). With our triple knockdown approach to deplete all three endophilin A1/A2/A3 proteins as performed in the original study, we observed decreased JEV entry corresponding to reduced virus replication and titres. We also observe an increase in JEV internalization in SH-SY5Y cells and in primary cortical neurons upon treatment with a specific inhibitor of GSK3β. This establishes that activation of an endophilin-mediated pathway can lead to an increase in JEV internalization, and highlights the crucial role of GSK3β in regulating virus endocytosis. Endophilin proteins are associated with the distinctive feature of membrane curvature and formation of long tubular structures for vesicular trafficking. Ultrastructural studies also corroborate JEV localization whithin longer cavities at the cell surface that are distinct from classical clathrin-coated vesicles. Through immunofluorescence, we also observe strong colocalization of virions with Endophilin, and a substantial increase in endophilin-positive structures or assemblies upon JEV internalization. This aligns with FEME, where activation of the pathway with its specific cargo or serum stimulation increases endophilin-positive structures. Overexpression of endophilin truncated mutant domains showed a reduction in virus entry and replication, with both the domains i.e., the cargo prolone-rich binding domain SH_3_, as well as the local membrane curvature sensing and formation, N-BAR domain being essential.

Endophilin mediated endocytosis has not been explored widely in the context of virus infection, with only one study showing that silencing of endophilin A2 decreases Enterovirus 71 infection (53). Endophilin has been shown to interact with Moloney murine leukemia virus (Mo-MuLV) Gag protein, and with the mouse cytomegalovirus pM50 protein, and functions in membrane curvature formation and virion production (54,55). Endophilin B2 has been shown to be essential for Influenza A viral RNA nuclear entry and replication (56).

Actin cytoskeleton network participates in both clathrin-dependent and independent pathways by providing a mechanical force to bend membrane and form endocytic pits. Multiple viruses such as CSFV, HIV-1, HCV, pseudorabies virus, HPV-16, Rhinovirus etc. utilize the actin machinery to facilitate their transport within the cell (57–62), A study from our laboratory performed an RNA interference-based screening of 136 human membrane trafficking genes in IMR-32 cells (human neuroblastoma) and identified RHOA, RAC1, PAK1, along with ARP2/3 complex, and N-WASP family proteins to be essential for JEV replication (17). As demonstrated in the previous study, pharmacological disruption of the actin polymerization and depolymerisation, or inhibitors of the ARP2/3 complex, and N-WASP led to a decrease in virus internalization. These results were also supported by immunofluorescence studies wherein remodelling of the actin cytoskeleton structures with dense F-actin clusters were observed in infected cells, as compared to the characteristic actin filaments in uninfected cells.

RTKs has been associated with internalization of multiple viruses. EGFR is a member of human ErbB family of RTKs with three characteristics domains: an extracellular ligand binding domain, a transmembrane, and a cytoplasmic domain. Receptor activation can be ligand-dependent as well as -independent, with the physiological role of the ligand-independent activation not completely understood (63–65). EGFR can be trafficked through clathrin dependent or independent pathways depending on the cellular growth conditions (66). The CIE of EGFR is observed upon stimulation with high EGF concentrations, as there is a rapid need for receptors to undergo degradation in order to maintain the cellular homeostasis (23,33,67). The receptor undergoes substantial ubiquitination at high ligand concentration mediated through an E3 ligase Cbl and hence switches to a CIE pathway (68,69).

Studies have shown that JEV internalization activates EGFR signaling through the EGFR-PI3K signaling axis promoting the RhoA-mediated F-actin polymerization in neuronal cells (19). The activation of the EGFR-ERK signaling cascade during the early time of infection was also related with suppression of the interferon response in hBMECs (70). However, it still remains unclear how EGFR interacts with JEV to facilitate entry. In our present study, we observed that virus entry also resulted in an activation of EGFR signaling as observed with an increase in the phosphorylated EGFR levels, while inhibition of EGFR signaling led to a decrease in JEV internalization. These results highlight the central role of intracellular kinase domain along with the activation of downstream signaling cascade for virus entry. These results were further confirmed by the genetic knockdown of EGFR, which led to a decrease in JEV internalization. A reduction in JEV attachment was also observed with EGFR depletion, which suggested the active role of the extracellular ligand-binding domain as well. Treatment of cells with EGFR extracellular domain binding antibody, cetuximab, and EGF led to a decrease in JEV attachment and internalization, further supporting the hypothesis that EGFR could potentially be involved in the initial binding of JEV with the host cell surface. Treatment of cells with EGF further showed a decrease in JEV binding at 4°c where endocytosis has not yet initiated. Hence, our data indicates that interference with the ligand binding domain of EGFR prevents JEV binding, and subsequently blocks virus internalization. Through high-resolution immunofluorescence microscopy, we observed strong colocalization of JEV envelope particles with EGFR at early times of infection, with the maximum colocalization observed at t=0 min, highlighting the possibility of a direct association between the two. Interestingly, through BLI we observed a strong interaction of JEV ED3 domain with EGFR, but not with the mutant isoform. Overall, these data indicate a pivotal role of the EGFR signaling and extracellular ligand binding domain to be essential for JEV endocytosis, and point towards its role as a specific receptor.

Our study describes two critical host proteins that are exploited by JEV for entry in neuronal cells. However, targeting these fundamental physiological signaling hubs for antiviral development is likely to be challenging, and highlights the need for a more nuanced understanding of virus-host interactions.

## Materials & Methods

### Ethics Statement

All animal experiments were approved by the Institutional Animal Ethics Committee of the Regional Centre for Biotechnology (RCB/IAEC/2022/114). Mice were maintained and experiments were performed as per the guidelines of the Committee for the Purpose of Control and Supervision of Experiments on Animals (CPCSEA), Government of India.

### Cell lines and virus

Human neuroblastoma cell line SH-SY5Y was obtained from ATCC. Vero and C6/36 cell lines were obtained from the cell repository of National Centre for Cell Sciences Pune, India. SH-SY5Y cells were cultured in HiGlutaXL^TM^ Dulbecco’s modified eagle medium (DMEM), C6/36 ells in Leibovitz’s L-15 Medium, Vero cells in Minimum essential medium Eagle (MEM) and HeLa cells in DMEM. All media for culturing was supplemented with 10% FBS and 1X penicillin-streptomycin-glutamine (PSG).

JEV Vellore strain P20778 (GenBank accession no. AF080251) was generated in insect cell line C6/36 in 2 % FBS with 1X penicillin-streptomycin-glutamine (PSG).

### Virus generation and purification

C6/36 cells were infected with JEV 0.1 MOI at a confluency of 70-80 % for 1 h and were supplemented with 2% FBS containing L-15 media. Supernatant containing the virus particles were harvested when ∼80 % of the cells showed cytopathic effect at approximately 72 h post infection (pi). Virus particles were collected by centrifugation at 1,000 × g for 15 min at 4°c. Virus concentration was done through multiple cycles of centrifugation of the collected supernatant with Amicon® Ultra Centrifugal Filter, 100 kDa MWCO at 1,000 × g for 15 min at 4°c. For virus purification, the virus containing supernatant was first concentrated through PEG precipitation, followed with purification over 20 % sucrose cushion through ultracentrifugation at 80,000 × g for 4 h at 4°c (16).

### Virus titration

Virus titre estimation was performed by plaque assays on Vero cells. Cells were plated in 12-well plate and were infected with 10-fold serially diluted virus supernatant for 1 hr at 37°c. Virus inoculum was removed post 1 h and cells were washed with PBS twice after which they were overlayed with low-melting-point agarose and 2X MEM in 1:1 ratio for 5 days at 37°c until the plaques became visible. Cells were then fixed in 3.75 % formaldehyde overnight and the agarose plug were removed followed with staining with 0.1 % crystal violet blue for plaque visualization. Titres were determining by counting the number of plaques in the lowest dilutions were they were visible for counting which were represented as plaque forming units (pfu/ml).

### Small interfering RNA (siRNA) depletion and virus assays

Cells seeded in 24-well plate were double transfected (day 1 and day 2) with 25 nM siRNA on each day with DharmaFECT 1 transfection reagent according to the manufacturers instruction. Cells were transfected for 72 h before virus infection assays. Knockdown efficiency was checked through qRT-PCR and western blotting. Virus entry assays were performed by allowing the virus 10 MOI to bind on ice for 1 h followed with removing the virus inoculum. Cells were shifted to 37°c to allow virus internalization for 1 h and were washed with 1X PBS twice followed with treatment with trypsin to remove extracellular bound virus particles. Cells were then washed with PBS and were harvested with TRIzol for RNA extraction and qRT-PCR. For virus entry assay under FEME conditions, cells were infected with virus 10 MOI for 1 h at 37°c followed with virus inoculum removal. Cells were washed with PBS and were either harvested immediately with trypsin treatment (0 hpi) or kept for another 1 h in complete media (1 hpi) followed with trypsin treatment to harvest cells with TRIzol. qRT-PCR was performed with primers against JEV envelope region to measure the absolute viral copy numbers. For virus binding or attachment studies, cells were infected with virus 10 MOI on ice for 1 h to allow virus particles to bind. Cells were washed with chilled PBS and were harvested with TRIzol for RNA isolation and qRT-PCR. For virus infection assays, cells were infected with JEV 1 MOI for 1 h and were washed with PBS. Cells were then kept in complete media for 24 h and were harvested with TRIzol. qRT-PCR was done to measure virus RNA levels. For immunofluorescence based virus entry assays, cells were infected with virus of 100 MOI on ice and were shifted to 37°c for different time intervals subsequently fixing it with 4 % PFA.

### RNA isolation and Quantitative Real Time (qRT)-PCR

RNA extraction was performed with TRIzol RNAiso Plus reagent using a phenol-chloroform based method. Briefly, cells were harvested with TRIzol and were homogenized by pipetting multiple times. 100 ul of chloroform was added per sample and was incubated for 2 mins at RT followed with centrifugation at 12000 rpm for 15 min at 4°c to allow phase separation. The clear aqueous phase was separated and 250 ul isopropanol was added. The samples were incubated at room temperature for 15 min to allow RNA precipitation and were centrifuged at 12000 rpm for 15 min at 4°c. The supernatant was discarded and the pellet was washed with chilled 75 % ethanol with centrifugation at 12000 rpm for 15 min at 4°c. Samples were air dried and were resuspended in nuclease free water by heating at 55°c for 10 min. cDNA preparation was done using Prime-script Transcription kit and real-time PCR was performed using Quant Studio 6 (Applied Biosystems). All the primer sequences used are mentioned in supplementary table S1.

### Western blotting

Cell lysis was performed in lysis buffer containing 150 mM NaCl, 1% Triton X-100, 50 mM Tris-HCl pH 7.5, with 1 mM PMSF, and protease inhibitor cocktail for 60 min on ice. Supernatant collected were used for protein quantification using BCA assay kit. 5X loading was added to the cell lysates and these were heated at 95°c for 10 min. Equal concentration of protein lysates were run on a SDS-PAGE gel and transferred to PVDF membrane for immunoblotting. Membranes were cut according to the size of the desired protein and were blocked in 5 % not fat milk in 1X PBS/TBS for 1 h at RT. Primary antibody incubation was done in the blocking solution at the desired antibody dilution at 4°c overnight. Membrane were washed 3 times with 1x PBST/TBST (1 % tween in PBS/TBS). Secondary HRP-conjugated antibody was diluted in milk and was incubated for 1 h at RT. Membrane was washed thrice with 1x PBST/TBST, and bands were revealed with chemiluminescence using HRP substrate.

### Transmission electron microscopy

SH-SY5Y cells were infected with JEV (MOI 200), incubated in ice for 2 hours and were fixed at 30 seconds post infection in 2.5% glutaraldehyde and 2 % PFA in 0.1M sodium cacodylate buffer, followed by post-fixation in 1% osmium tetroxide, dehydration in ethanol, and embedded in LR white resin (Ted Pella). Blocks were sectioned using Leica UC7 Ultramicrotome 5-glass knives. Sections were examined at RT with a transmission electron microscope (Talos, L120C) at 11kx.

### Gene cloning and stable mutant cell line generation

Full-length (FL) Human Endophilin A2 (Endo-A2)/SH3GL1(Gene ID: 6455, NP_003016.1) and different functional domain mutants were cloned into mammalian expression vector pEGFP_C1 to obtain N-terminus GFP-tagged fusion protein. Domain specific regions were classified based on the NCBI database. Briefly, full length Endo-A2 (368 aa) was generated by PCR as BglII/EcoRI fragments (FL-F and FL-R) and subcloned into pEGFP_C1 at the same sites. Similarly, fragments expressing 22-368 aa and 1-308 aa were PCR amplified from cDNA, using primers (ΔH°-F and R and ΔSH_3_-F and R, respectively) with 5’ and 3’ tail containing BglII and EcoRI sites, for directional cloning into the pEGFP_C1 vector. These constructs were referred to as EGFP-ΔHO (22-368 aa) and EGFP-ΔSH3 (1-308 aa), respectively.

Overlap extension PCR was used to construct the BAR-domain mutant. This chimera consists of N-terminal 1-24 residues fused to the C-terminal 248-368 aa of Endo A2. In the first round PCR, fragment 1 viz. 1-24 aa was amplified using (ΔH°-F and ΔBAR_1-R) primers whereas fragment 2 viz. 248-368 aa was amplified using the primers (ΔBAR_2-F and ΔSH_3_-R), and both the fragments were then gel purified. The chimeric primers viz. ΔBAR_2-F and ΔBAR_1-R contains ∼12 bp overhangs at the 5’ end, which are complementary to fragment 1 and fragment 2, respectively. In a 2-step overlap extension, two purified fragments (fragment 1 and 2) were mixed in equimolar concentration (0.1 pmol each) for overlap extension PCR performed as follows: thirteen cycles of 10 sec at 92°c (denaturation), 1 min at 60°c (annealing) and 72°c for 50 sec (extension). In step 2, resulting PCR product was taken as a template for the nested PCR, where extreme end primers viz. ΔBAR_1-F and ΔBAR_2-R were used for the amplification of the full length (147aa) hybrid fragment. The fusion product was then digested and ligated into pEGFP_C1 vector at BglII-EcoRI sites to generate EGFP-ΔBAR. The recombinant plasmids were transformed into DH5α E. coli strains and screened for the positive clones. All constructs were verified by sequencing. To generate the stable cell lines expressing different domain mutants of Endo-A2, SH-SY5Y cells were seeded in 24 well plate overnight and transfected with the respective plasmids viz. pEGFP_C1 (empty vector), EGFP-FL (full length), EGFP-ΔHO, EGFP-ΔSH3 and EGFP-ΔBAR. Briefly, cells were transfected with plasmid concentration of 2 µg using Lipofectamine™ 3000 Transfection Reagent according to the manufacturer’s protocol. Stably expressing cells with EGFPC1 constructs were selected with neomycin (800 µg/ml) for 4-weeks and were sorted using flow cytometry for varied GFP intensities. Cells with high GFP signals were grown in antibiotic selection pressure and expression of tagged-protein variants was confirmed by immunofluorescence and immunoblotting. All the primer sequences used for cloning are mentioned in supplementary table S2.

### Primary cortical neurons isolation and culture

Primary cortical neurons were isolated from a previously described protocol (71). Briefly,. embryos were collected from pregnant mice at embryonic day E.16.5 from C57/BL6 mice through decapitation from the pregnant mice in ice-cold dissociation media, HBSS (1X sodium pyruvate, 20% glucose, 1 M HEPES, pH 7.3). The cortices were gently dissected from the brain and were collected in the dissociation media. Tissues were washed and digested with trypsin and DNAse I at 37°c for 20 min. Single cell suspension were collected, washed twice and finally resuspended in neurobasal medium supplemented with 10% FBS, 20% glucose, 1X Sodium pyruvate, and antibiotics. The neurons isolated were plated on poly-l-lysine coated plates. The media was changed the following day upon adherence of the neurons on the plate with maintenance media (neurobasal B-27, 1X glutamine, penicillin-streptomycin solution).

### Antibody inhibition assays

Cells were seeded in 96-well plate and were allowed to bind on ice for 1 h with 50 ug/ml of the control (negative) mouse IgG and with specific EGFR antibody against the epitope binding domain (sc-120). This was followed with chilled ice cold PBS washes twice and were subsequently infected with JEV (1 MOI) for 30 min on ice. Cells were washed with chilled PBS and were incubated with complete media for 24 h. Cells were harvested post 24 h and were proceeded with RNA extraction and qRT-PCR. Plaque assays was performed to check virus titres collected from the supernatant. For virus binding/entry assays, post incubation with antibodies, cells were infected with JEV (10 MOI) for 30 min on ice and were washed with chilled PBS twice before harvesting with TRIzol (0 hpi) or adding complete media for 1 h at 37°c (1 hpi). Cells were washed and were treated with trypsin to remove non-internalized virus particles followed with PBS wash and were harvested with TRIzol reagent for RNA extraction and qRT-PCR.

### EGF stimulation and virus infection assay

Cells were seeded in 24-well plates overnight and were stimulated with 20 % serum for 15 min at 37°c to activate FEME. For virus binding assay, cells were either EGF stimulated or not, followed with incubation with both virus (10 MOI) and increasing concentration of EGF at 4°c for 1 h. Cells were harvested followed with RNA extraction, cDNA preparation and qRT-PCR. For entry assay, cells were stimulated with increasing concentration of EGF for 15 mins at 37°c and were subsequently allowed to bind with JEV (10 MOI) and with the increasing EGF concentration on ice for 1 h. Cells were washed with ice cold PBS and were incubated at 37°c for 1 h with complete media. Cells were harvested with TRIzol post 1 h after trypsin treatment. For studies of virus replication, the serum and EGF stimulated cells were infected with 1 MOI virus followed with harvesting at 24 hpi.

### Cargo uptake assay

For endocytic cargo uptake assays, cells were seeded in coverslip coated 24-well plate overnight. Post-transfection with siRNA for 72 h, for Tf uptake assay, cells were serum starved for 30 mins at 37°c. Cells were pulsed with Alexa fluor labelled Tf of concentration 20 ng/well for 5 min at 37°c. Cells were washed with acid wash buffer and subsequently with PBS twice to remove uninternalized Tf. Cells were fixed with 4 % PFA for 15 min at 37°c. For EGF uptake assay, cells were never serum starved, and were pulsed with different EGF concentrations of 10, 20, 50, or 180 ng/ml for 5 min at 37°c. Cells were then washed with PBS twice to remove uninternalized EGF and were subsequently fixed with 4 % PFA for 15 min at 37°c.

### Cell viability assay

SH-SY5Y cells were seeded in 96-well plate and were either left untreated (DMSO control) or treated with the drugs with the different concentrations. After 5 h post treatment, MTT was added at a final concentration of 0.5 mg/ml for 3 h followed by addition of solubilization solution to stop the reaction. OD was measured at 570 nm using a plate reader. The percentage cell viability was then calculated and normalized to DMSO treated control cells.

## Immunofluorescence studies

### Immunofluorescence staining and image acquisition

Cells grown on coverslips were either transfected with siRNA, plasmid or were infected with JEV MOI 100 for immunofluorescence based entry experiments. Virus infection was performed on ice for 1 h followed with subsequent removal of the inoculum, washing and incubating the cells at different time intervals at 37°c. Cells were then fixed with 4 % PFA for 15 min at 37°c. Immunostaining was performed by permeabilizing with 0.3 % tween or 0.05 % saponin, blocking in 1 % BSA in PBS, incubating in primary antibody for 1 h at RT or overnight at 4°c, followed by fluorescent tagged secondary antibody for 1 h at RT. Cells were mounted under Prolong gold reagent with DAPI. Cells were imaged using Laser confocal scanning Leica TCS SP8 microscope or Elyra PS1 (Carl Zeiss Super-resolution microscope) using 63 x objectives.

### Phalloidin staining

Cells grown on coverslips were fixed in 4 % PFA solution for 15 min at room temperature and were washed thrice with PBS. Cells were permeabilized in 0.3 % tween for 15 mins, washed and blocked in 1 % BSA in PBS. Cells were incubated with primary antibody for 1 h at RT or overnight at 4°c, followed by fluorescent tagged secondary antibody combined with Alexa Fluor 546-conjugated phalloidin (1:40, Invitrogen, A22283) for 1 h at RT. Cells were washed, stained with DAPI and mounted in Prolong gold reagent.

### Quantification of cargo uptake assay images

Image quantification of cargo uptake assays was performed using Image J software. Briefly, approximately ∼100 cells per coverslip were used for analysis. Bright field image was used to draw cell boundary to generate region of interest against each cell with manual free hand tool. This was followed with integrated fluorescent intensity calculation with measure tool from respective channels. Graph were plotted as mean ± SEM.

### Quantification of colocalization for SIM images

For quantification of colocalization in SIM images, an object-object based colocalization approach was utilized in Imaris version 9.9. A region of interest was created around each cell using the surface function. Spots were generated for each channel based on the size of the puncta. Briefly, the size of few puncta were measured, and subsequently based on the size of the spots, a size was set for each channel. The threshold was set manually for each cell for all the channels to obtain all the spots visible in three-dimensional viewer. The spots for all the channels to be quantified were created similarly. The colocalization between any of the channels was measured with calculating the shortest distance between two spots. A shortest distance parameter of 0.5 um was set. Based on this the spots with distance within 0.5 um were considered as colocalized.

### JEV ED3 protein purification

JEV ED3 protein purification was performed from a previously described protocol from (10). Briefly, E. coli BL-21(DE3) cells transformed with pET28a plasmid carrying the ED3 cDNA (aa 303 to 398) with a c-terminal His-tag were grown at 37°c till the O.D_600_ reaches 0.6-0.8. The protein expression was induced by adding 1 mM IPTG (Isopropyl β-D-1-thiogalactopyranoside) at 30°C for 4h with constant shaking at 200 rpm. The induced bacterial pellet was lysed under denaturing conditions in lysis buffer for 1h with continuous swirling at 37°C (100 mM NaH PO, 10 mM Tris-HCl, 8 M urea, pH 8.0, 1:50 PI and 1:50 PMSF). Ni-NTA beads were added to the cell supernatant for protein purification and was eluted in 8 M urea containing 200mM imidazole at pH 8. The elute was dialyzed against 8 M, 6 M, 4 M, 3 M, 2 M, 1M urea and finally in PBS. Dialyzed protein was collected and centrifuged at 12000 rpm for 90 min at 4°c and supernatant was concentrated using 3 KDa Amicon filter. The protein concentration was determined using BCA and ED3 protein bands were confirmed using western blots as a single band at ∼10 KDa molecular weight.

### Biolayer Interferometry (BLI)

All the binding experiments were performed using an Octet RED96 (Forte Bio) at 30°C with shaking at 800 rpm. Prior to use, the Ni-NTA biosensors were loaded into the columns of a biosensor holding plate and pre-hydrated in assay buffer for 15 min. Protein JEV-ED3 with a c-terminal His-tag was immobilized on Ni-NTA biosensor tips (Forte Bio) at a concentration of 20 μg/mL in an assay buffer containing PBS. Binding interactions were measured by submerging the JEV-ED3 loaded biosensors in solutions containing different concentrations of recombinant Human EGFR Protein (ECD, hFc Tag) or (Isoform Viii, hFc Tag) in a two-fold dilution series. The association and dissociation were measured for 400 s. The biosensors were regenerated for 3-4 cycles using a 10 mM glycine solution, pH 2.0, and then neutralized in the assay buffer. After regeneration, the biosensors were recharged using a 10 mM NiCl_2_ solution for 60 s. The sensor with 0 μg/mL recombinant EGFR was used for reference subtraction. The binding data were fitted using a 2:1 (Heterologous binding model) binding model using Octet Data Analysis software to evaluate the equilibrium dissociation constants (*K_D_*).

### Statistical analysis and software

GraphPad Prism version 9 was used to perform all the statistical analysis. Statistical analysis was done using paired/unpaired Student’s t-test, Kruskal-Wallis test with Dunn’s multiple comparisons test or one-way ANOVA followed by Dunnett’s/Šídák’s multiple comparisons multiple comparison test. Error bar indicates means ± SD/SEM. Microscopic image analysis was performed on Image J or Imaris version 9.9.

## Acknowledgements

All Virology lab members are acknowledged for useful discussions and support.

## References

1. Kalia M, Jameel S. Virus entry paradigms. Amino Acids [Internet]. 2011 Nov [cited 2024 Dec 5];41(5):1147–57. Available from: https://pubmed.ncbi.nlm.nih.gov/19826903/

2. Helenius A. Virus Entry: Looking Back and Moving Forward. J Mol Biol. 2018 Jun;430(13):1853–62.

3. McMahon HT, Boucrot E. Molecular mechanism and physiological functions of clathrin-mediated endocytosis. Nat Rev Mol Cell Biol. 2011 Jul 22;12(8):517–33.

4. Briant K, Redlingshöfer L, Brodsky FM. Clathrin’s life beyond 40: Connecting biochemistry with physiology and disease. Curr Opin Cell Biol. 2020 Aug;65:141–9.

5. Ferreira APA, Boucrot E. Mechanisms of Carrier Formation during Clathrin-Independent Endocytosis. Trends Cell Biol. 2018 Mar;28(3):188–200.

6. Renard HF, Boucrot E. Unconventional endocytic mechanisms. Curr Opin Cell Biol. 2021 Aug;71:120–9.

7. Thottacherry JJ, Sathe M, Prabhakara C, Mayor S. Spoiled for Choice: Diverse Endocytic Pathways Function at the Cell Surface. Annu Rev Cell Dev Biol. 2019 Oct 6;35(1):55–84.

8. Turtle L, Solomon T. Japanese encephalitis — the prospects for new treatments. Nature Reviews Neurology 2018 14:5 [Internet]. 2018 Apr 26 [cited 2024 Dec 9];14(5):298–313. Available from: https://www.nature.com/articles/nrneurol.2018.30

9. Sharma KB, Vrati S, Kalia M. Pathobiology of Japanese encephalitis virus infection. Mol Aspects Med [Internet]. 2021 Oct 1 [cited 2024 Dec 5];81. Available from: https://pubmed.ncbi.nlm.nih.gov/34274157/

10. Nain M, Mukherjee S, Karmakar SP, Paton AW, Paton JC, Abdin MZ, et al. GRP78 Is an Important Host Factor for Japanese Encephalitis Virus Entry and Replication in Mammalian Cells. J Virol. 2017 Mar 15;91(6).

11. Niu J, Jiang Y, Xu H, Zhao C, Zhou G, Chen P, et al. TIM-1 Promotes Japanese Encephalitis Virus Entry and Infection. Viruses. 2018 Nov 14;10(11):630.

12. Chien YJ, Chen WJ, Hsu WL, Chiou SS. Bovine lactoferrin inhibits Japanese encephalitis virus by binding to heparan sulfate and receptor for low density lipoprotein. Virology. 2008 Sep;379(1):143–51.

13. Chiou S, Liu H, Chuang C, Lin C, Chen W. Fitness of Japanese encephalitis virus to Neuro-2a cells is determined by interactions of the viral envelope protein with highly sulfated glycosaminoglycans on the cell surface. J Med Virol. 2005 Aug 23;76(4):583– 92.

14. Nain M, Abdin MZ, Kalia M, Vrati S. Japanese encephalitis virus invasion of cell: allies and alleys. Reviews in Medical Virology. John Wiley and Sons Ltd; 2016. p. 129–41.

15. Liang JJ, Yu CY, Liao CL, Lin YL. Vimentin binding is critical for infection by the virulent strain of Japanese encephalitis virus. Cell Microbiol. 2011 Sep;13(9):1358–70.

16. Kalia M, Khasa R, Sharma M, Nain M, Vrati S. Japanese Encephalitis Virus Infects Neuronal Cells through a Clathrin-Independent Endocytic Mechanism. J Virol. 2013 Jan;87(1):148–62.

17. Khasa R, Sharma P, Vaidya A, Vrati S, Kalia M. Proteins involved in actin filament organization are key host factors for Japanese encephalitis virus life-cycle in human neuronal cells. Microb Pathog. 2020 Dec;149:104565.

18. Zhu YZ, Xu QQ, Wu DG, Ren H, Zhao P, Lao WG, et al. Japanese Encephalitis Virus Enters Rat Neuroblastoma Cells via a pH-Dependent, Dynamin and Caveola-Mediated Endocytosis Pathway. J Virol. 2012 Dec 15;86(24):13407–22.

19. Xu Q, Cao M, Song H, Chen S, Qian X, Zhao P, et al. Caveolin-1-Mediated Japanese Encephalitis Virus entry Requires a Two-Step Regulation of Actin Reorganization. Future Microbiol. 2016 Oct 17;11(10):1227–48.

20. Khasa R, Vaidya A, Vrati S, Kalia M. Membrane trafficking RNA interference screen identifies a crucial role of the clathrin endocytic pathway and ARP2/3 complex for Japanese encephalitis virus infection in HeLa cells. Journal of General Virology. 2019 Feb 1;100(2):176–86.

21. Liu CC, Zhang YN, Li ZY, Hou JX, Zhou J, Kan L, et al. Rab5 and Rab11 Are Required for Clathrin-Dependent Endocytosis of Japanese Encephalitis Virus in BHK-21 Cells. J Virol. 2017 Oct;91(19).

22. Yang S, He M, Liu X, Li X, Fan B, Zhao S. Japanese encephalitis virus infects porcine kidney epithelial PK15 cells via clathrin- and cholesterol-dependent endocytosis. Virol J. 2013 Dec 12;10(1):258.

23. Boucrot E, Ferreira APA, Almeida-Souza L, Debard S, Vallis Y, Howard G, et al. Endophilin marks and controls a clathrin-independent endocytic pathway. Nature. 2015 Jan 22;517(7535):460–5.

24. Casamento A, Boucrot E. Molecular mechanism of Fast Endophilin-Mediated Endocytosis. Biochemical Journal. 2020 Jun 26;477(12):2327–45.

25. Carlin CR. Role of EGF Receptor Regulatory Networks in the Host Response to Viral Infections. Front Cell Infect Microbiol. 2022 Jan 10;11.

26. Noh SS, Shin HJ. Role of Virus-Induced EGFR Trafficking in Proviral Functions. Biomolecules. 2023 Dec 9;13(12):1766.

27. Tanaka T, Zhou Y, Ozawa T, Okizono R, Banba A, Yamamura T, et al. Ligand-activated epidermal growth factor receptor (EGFR) signaling governs endocytic trafficking of unliganded receptor monomers by non-canonical phosphorylation. Journal of Biological Chemistry. 2018 Feb;293(7):2288–301.

28. Sigismund S, Woelk T, Puri C, Maspero E, Tacchetti C, Transidico P, et al. Clathrin-independent endocytosis of ubiquitinated cargos. Proceedings of the National Academy of Sciences. 2005 Feb 22;102(8):2760–5.

29. Macia E, Ehrlich M, Massol R, Boucrot E, Brunner C, Kirchhausen T. Dynasore, a Cell-Permeable Inhibitor of Dynamin. Dev Cell. 2006 Jun;10(6):839–50.

30. Kjaerulff O, Brodin L, Jung A. The Structure and Function of Endophilin Proteins. Cell Biochem Biophys. 2011 Jul 25;60(3):137–54.

31. Sigismund S, Woelk T, Puri C, Maspero E, Tacchetti C, Transidico P, et al. Clathrin-independent endocytosis of ubiquitinated cargos. Proc Natl Acad Sci U S A. 2005 Feb 22;102(8):2760–5.

32. Sigismund S, Argenzio E, Tosoni D, Cavallaro E, Polo S, Di Fiore PP. Clathrin-Mediated Internalization Is Essential for Sustained EGFR Signaling but Dispensable for Degradation. Dev Cell. 2008 Aug;15(2):209–19.

33. Caldieri G, Barbieri E, Nappo G, Raimondi A, Bonora M, Conte A, et al. Reticulon 3– dependent ER-PM contact sites control EGFR nonclathrin endocytosis. Science (1979). 2017 May 12;356(6338):617–24.

34. Ferreira APA, Casamento A, Carrillo Roas S, Halff EF, Panambalana J, Subramaniam S, et al. Cdk5 and GSK3β inhibit fast endophilin-mediated endocytosis. Nat Commun. 2021 Apr 23;12(1):2424.

35. Spector I, Shochet NR, Kashman Y, Groweiss A. Latrunculins: Novel Marine Toxins That Disrupt Microfilament Organization in Cultured Cells. Science (1979). 1983 Feb 4;219(4584):493–5.

36. Bubb MR, Senderowicz AM, Sausville EA, Duncan KL, Korn ED. Jasplakinolide, a cytotoxic natural product, induces actin polymerization and competitively inhibits the binding of phalloidin to F-actin. Journal of Biological Chemistry. 1994 May;269(21):14869–71.

37. Goddette DW, Frieden C. Actin polymerization. The mechanism of action of cytochalasin D. Journal of Biological Chemistry. 1986 Dec;261(34):15974–80.

38. Shoji K, Ohashi K, Sampei K, Oikawa M, Mizuno K. Cytochalasin D acts as an inhibitor of the actin–cofilin interaction. Biochem Biophys Res Commun. 2012 Jul;424(1):52–7.

39. Guerriero CJ, Weisz OA. N-WASP inhibitor wiskostatin nonselectively perturbs membrane transport by decreasing cellular ATP levels. American Journal of Physiology-Cell Physiology. 2007 Apr;292(4):C1562–6.

40. Yang L, Xu J, Guo L, Guo T, Zhang L, Feng L, et al. Porcine Epidemic Diarrhea Virus-Induced Epidermal Growth Factor Receptor Activation Impairs the Antiviral Activity of Type I Interferon. J Virol. 2018 Apr 15;92(8).

41. Sabino C, Bender D, Herrlein ML, Hildt E. The Epidermal Growth Factor Receptor Is a Relevant Host Factor in the Early Stages of The Zika Virus Life Cycle *In Vitro*. J Virol. 2021 Sep 27;95(20).

42. Wang Q, Pan W, Wang S, Pan C, Ning H, Huang S, et al. Protein Tyrosine Phosphatase SHP2 Suppresses Host Innate Immunity against Influenza A Virus by Regulating EGFR-Mediated Signaling. J Virol. 2021 Feb 24;95(6).

43. Li S, Schmitz KR, Jeffrey PD, Wiltzius JJW, Kussie P, Ferguson KM. Structural basis for inhibition of the epidermal growth factor receptor by cetuximab. Cancer Cell. 2005 Apr;7(4):301–11.

44. Sorkin A. Internalization of the epidermal growth factor receptor: role in signalling. Biochem Soc Trans. 2001 Aug 1;29(4):480–4.

45. Luca VC, AbiMansour J, Nelson CA, Fremont DH. Crystal Structure of the Japanese Encephalitis Virus Envelope Protein. J Virol. 2012 Feb 15;86(4):2337–46.

46. Thongtan T, Wikan N, Wintachai P, Rattanarungsan C, Srisomsap C, Cheepsunthorn P, et al. Characterization of putative Japanese encephalitis virus receptor molecules on microglial cells. J Med Virol. 2012 Apr 15;84(4):615–23.

47. Das S, Ravi V, Desai A. Japanese encephalitis virus interacts with vimentin to facilitate its entry into porcine kidney cell line. Virus Res. 2011 Sep;160(1–2):404–8.

48. Das S, Laxminarayana SV, Chandra N, Ravi V, Desai A. Heat shock protein 70 on Neuro2a cells is a putative receptor for Japanese encephalitis virus. Virology. 2009 Mar;385(1):47–57.

49. Ren J, Ding T, Zhang W, Song J, Ma W. Does Japanese encephalitis virus share the same cellular receptor with other mosquito-borne flaviviruses on the C6/36 mosquito cells? Virol J. 2007 Dec 6;4(1):83.

50. Das S, Chakraborty S, Basu A. Critical role of lipid rafts in virus entry and activation of phosphoinositide 31 kinase/Akt signaling during early stages of Japanese encephalitis virus infection in neural stem/progenitor cells. J Neurochem. 2010 Oct 31;115(2):537–49.

51. Chuang CK, Yang TH, Chen TH, Yang CF, Chen WJ. Heat shock cognate protein 70 isoform D is required for clathrin-dependent endocytosis of Japanese encephalitis virus in C6/36 cells. Journal of General Virology [Internet]. 2015 Apr 1 [cited 2024 Dec 3];96(4):793–803. Available from: https://www.microbiologyresearch.org/content/journal/jgv/10.1099/jgv.0.000015

52. Zhu YZ, Xu QQ, Wu DG, Ren H, Zhao P, Lao WG, et al. Japanese Encephalitis Virus Enters Rat Neuroblastoma Cells via a pH-Dependent, Dynamin and Caveola-Mediated Endocytosis Pathway. J Virol. 2012 Dec 15;86(24):13407–22.

53. Chen SL, Liu YG, Zhou YT, Zhao P, Ren H, Xiao M, et al. Endophilin-A2-mediated endocytic pathway is critical for enterovirus 71 entry into caco-2 cells. Emerg Microbes Infect. 2019 Jan 28;8(1):773–86.

54. Wang MQ, Kim W, Gao G, Torrey TA, Morse HC, De Camilli P, et al. Endophilins interact with Moloney murine leukemia virus Gag and modulate virion production. J Biol. 2003 Dec 4;3(1):4.

55. Lemnitzer F, Raschbichler V, Kolodziejczak D, Israel L, Imhof A, Bailer SM, et al. Mouse cytomegalovirus egress protein pM50 interacts with cellular endophilin-A2. Cell Microbiol. 2013 Feb;15(2):335–51.

56. Serfass JM, Takahashi Y, Zhou Z, Kawasawa YI, Liu Y, Tsotakos N, et al. Endophilin B2 facilitates endosome maturation in response to growth factor stimulation, autophagy induction, and influenza A virus infection. Journal of Biological Chemistry. 2017 Jun;292(24):10097–111.

57. Schelhaas M, Ewers H, Rajamäki ML, Day PM, Schiller JT, Helenius A. Human Papillomavirus Type 16 Entry: Retrograde Cell Surface Transport along Actin-Rich Protrusions. PLoS Pathog. 2008 Sep 5;4(9):e1000148.

58. Li XM, Xu K, Wang JY, Guo JY, Wang XH, Zeng L, et al. The actin cytoskeleton is important for pseudorabies virus infection. Virology. 2024 Dec;600:110233.

59. Baktash Y, Madhav A, Coller KE, Randall G. Single Particle Imaging of Polarized Hepatoma Organoids upon Hepatitis C Virus Infection Reveals an Ordered and Sequential Entry Process. Cell Host Microbe. 2018 Mar;23(3):382–394.e5.

60. Serrano T, Frémont S, Echard A. Get in and get out: Remodeling of the cellular actin cytoskeleton upon HIV-1 infection. Biol Cell. 2023 Apr 15;115(4).

61. Cheng Y, Lou J xiu, Liu C chun, Liu Y yun, Chen X nan, Liang X dong, et al. Microfilaments and Microtubules Alternately Coordinate the Multistep Endosomal Trafficking of Classical Swine Fever Virus. J Virol. 2021 Apr 26;95(10).

62. Jubrail J, Africano-Gomez K, Herit F, Mularski A, Bourdoncle P, Oberg L, et al. Arpin is critical for phagocytosis in macrophages and is targeted by human rhinovirus. EMBO Rep. 2020 Jan 7;21(1).

63. Ceresa BP, Peterson JL. Cell and Molecular Biology of Epidermal Growth Factor Receptor. In 2014. p. 145–78.

64. Singh B, Carpenter G, Coffey RJ. EGF receptor ligands: recent advances. F1000Res. 2016 Sep 8;5:2270.

65. Byrne PO, Hristova K, Leahy DJ. EGFR forms ligand-independent oligomers that are distinct from the active state. Journal of Biological Chemistry. 2020 Sep;295(38):13353–62.

66. Tomas A, Futter CE, Eden ER. EGF receptor trafficking: consequences for signaling and cancer. Trends Cell Biol. 2014 Jan;24(1):26–34.

67. Watanabe S, Boucrot E. Fast and ultrafast endocytosis. Curr Opin Cell Biol. 2017 Aug;47:64–71.

68. Sigismund S, Woelk T, Puri C, Maspero E, Tacchetti C, Transidico P, et al. Clathrin-independent endocytosis of ubiquitinated cargos. Proceedings of the National Academy of Sciences. 2005 Feb 22;102(8):2760–5.

69. Sigismund S, Algisi V, Nappo G, Conte A, Pascolutti R, Cuomo A, et al. Threshold-controlled ubiquitination of the EGFR directs receptor fate. EMBO J. 2013 Jun 25;32(15):2140–57.

70. Zhang YG, Chen HW, Zhang HX, Wang K, Su J, Chen YR, et al. EGFR Activation Impairs Antiviral Activity of Interferon Signaling in Brain Microvascular Endothelial Cells During Japanese Encephalitis Virus Infection. Front Microbiol. 2022 Jun 30;13.

71. Prajapat SK, Mishra L, Khera S, Owusu SD, Ahuja K, Sharma P, et al. Methotrimeprazine is a neuroprotective antiviral in JEV infection via adaptive ER stress and autophagy. EMBO Mol Med. 2024 Jan 2;16(1):185–217.

