## Supplementary material for "Endophilin mediated endocytosis and Epidermal growth factor receptor govern Japanese encephalitis virus entry and infection in neuronal cells": Supplemantary Data

**Supplementary data**

Supplementary figure S1-S10

Supplementary table S1-S3

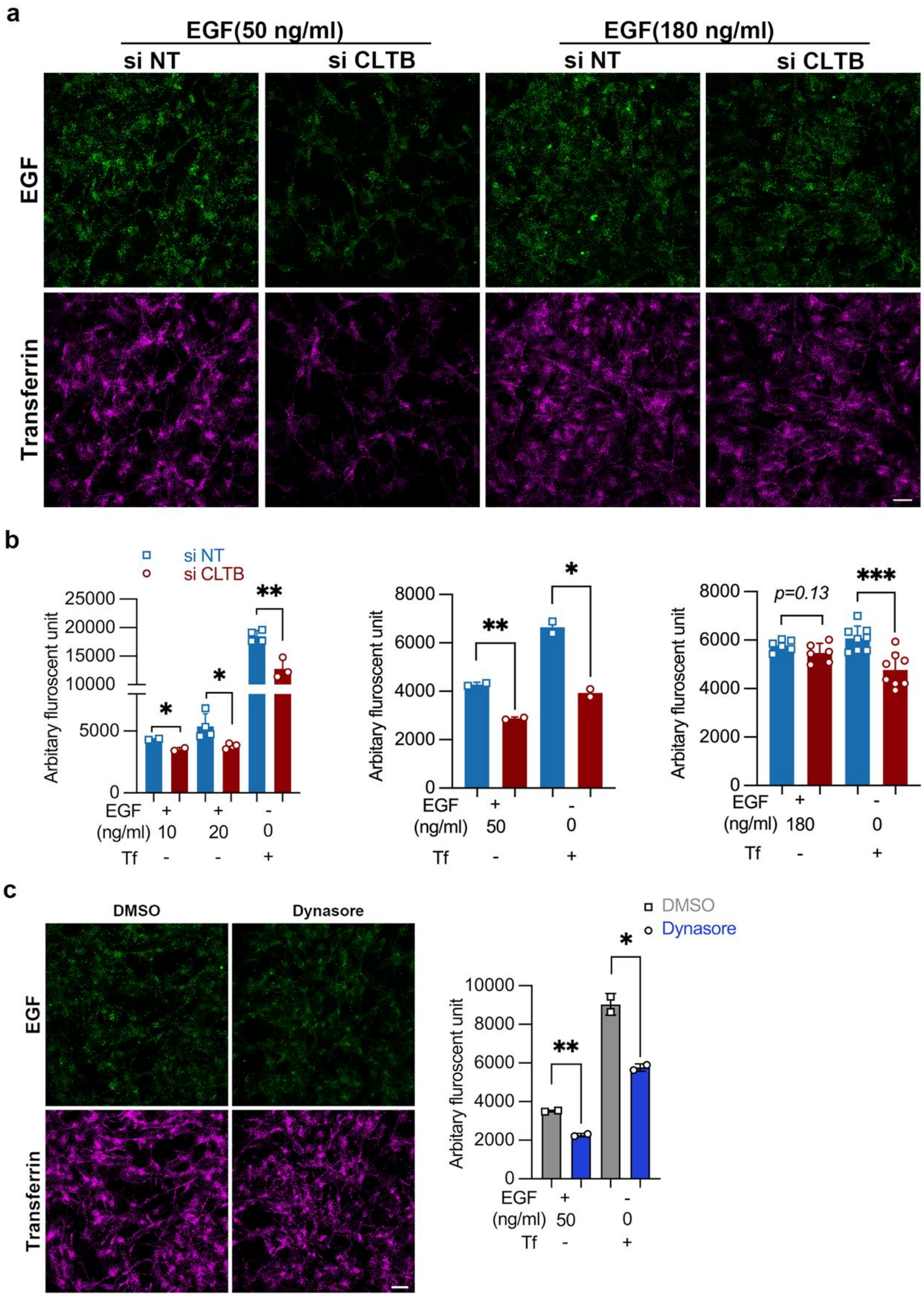


**Figure S1: Effect of CLTB knockdown on cargo internalization.** (a-b) SH-SY5Y cells transfected with siNT and siCLTB (72 h), were treated low (10, 20, 50 ng/ml), and high (180 ng/ml) concentration of Alexa fluor 555 EGF along with Alexa fluor 647 Tf (20 ng/ml) for 5 min at 37$^{O}$c. (a) Representative images indicating cargo uptake. Images were acquired using 63x objective, Scale: 20 µm. (b) Quantification of cargo uptake with a bar graph representing the total fluorescent intensities from 2 or more independent coverslips (~ 100 cells per coverslip).

(c) SH-SY5Y cells seeded in Nu-serum containing media were pre-treated with either DMSO or 80 µM of dynasore for 1 h at $37^{O}$c. Cells were then given a pulse of Alexa fluor 555 EGF (50 ng/ml) and Alexa fluor 647 Tf (20 ng/well) for 5 min at $37^{O}$c. Images show uptake of EGF and Tf upon treatment with either DMSO/dynasore. Scale, 20 µm. Bar graph compares the effect of dynasore treatment with DMSO control on the fluorescent intensities of EGF and Tf cargo. Total fluorescent intensities were quantified using Image J software with ~100 cells/coverslip. Individual values are representative of mean ± S.E.M. Statistical analysis was determined with student's unpaired two-tailed t-test, NEJM: 0.12 (ns), 0.033 (*), 0.002(**), <0.001(***).

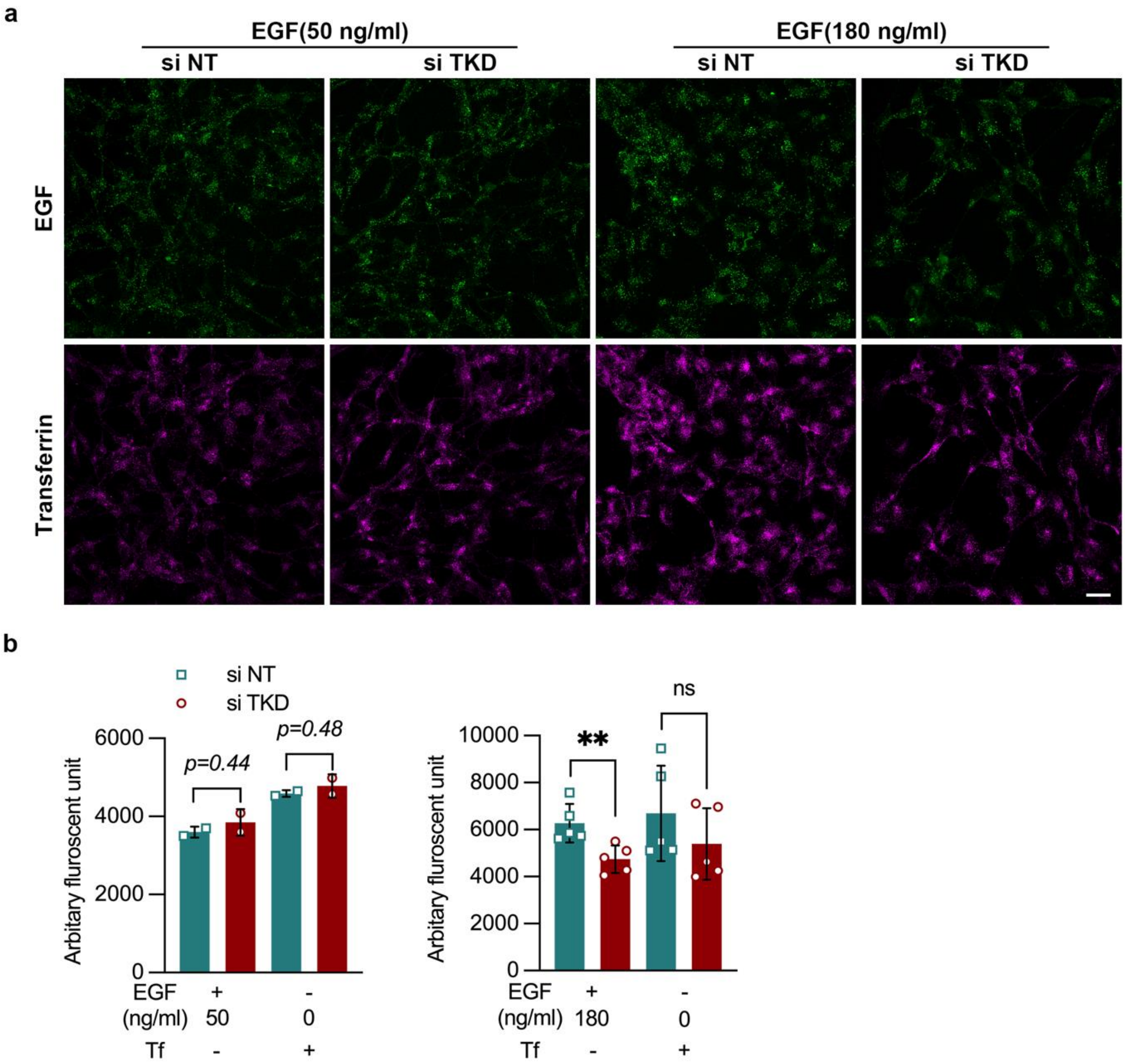


**Figure S2: Effect of Endophilin TKD on Tf and EGF uptake.** SH-SY5Y cells transfected with siNT and siTKD (72 h), were treated with Alexa fluor 555 EGF (50 & 180 ng/ml) and Alexa fluor 647 Tf (20 ng/well) for 5 min at $37^{O}$c. (a) Representative images indicating cargo uptake. Scale, 20 µm. (b) The bar graph quantifies the total fluorescent units of EGF and Tf uptake under siNT and siTKD conditions. Analysis was performed using image J software with ~100 cells/coverslip, represented as mean ± S.E.M. Statistical analysis was determined with unpaired student's t-test, NEJM: 0.12 (ns),0.033 (*), 0.002(**), <0.001(***).

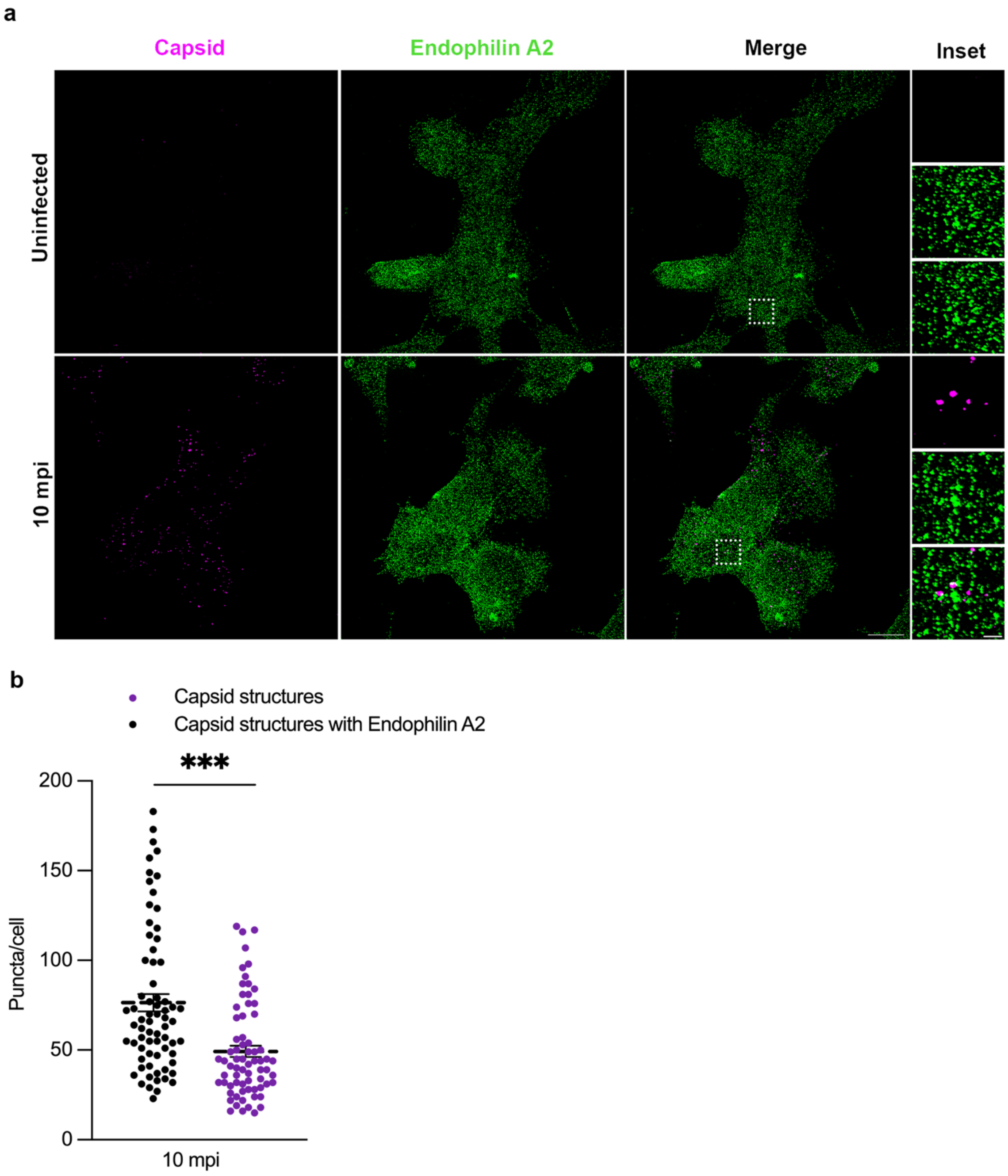


**Figure S3: JEV capsid structures colocalize with endophilin A2.** (a) SH-SY5Y cells were allowed to bind with 100 MOI virus on ice for 1 h and were subsequently shifted to $37^{O}$c for 10 min. Cells were immunostained for capsid (magenta) and endophilin A2 (green) and imaged on Elyra PS1 (Carl Zeiss Super-resolution microscope). Insets shows the magnified area from the confocal micrographs depicting the colocalization of endophilin A2 with capsid structures. Images are representative of two independent experiments with two coverslips for each experiment. Scale:10 µm, inset:1 µm. (b) The bar graph represents the quantification of the total number of capsid positive structures and the total number of capsid structures colocalized with endophilin A2 per cell (n>20 from two independent experiments). All values are represented as mean ± S.E.M; statistical analysis was determined with Mann Whitney test with 95% confidence level. Statistical significance: NEJM: 0.12 (ns),0.033 (*), 0.002(**), <0.001(***).

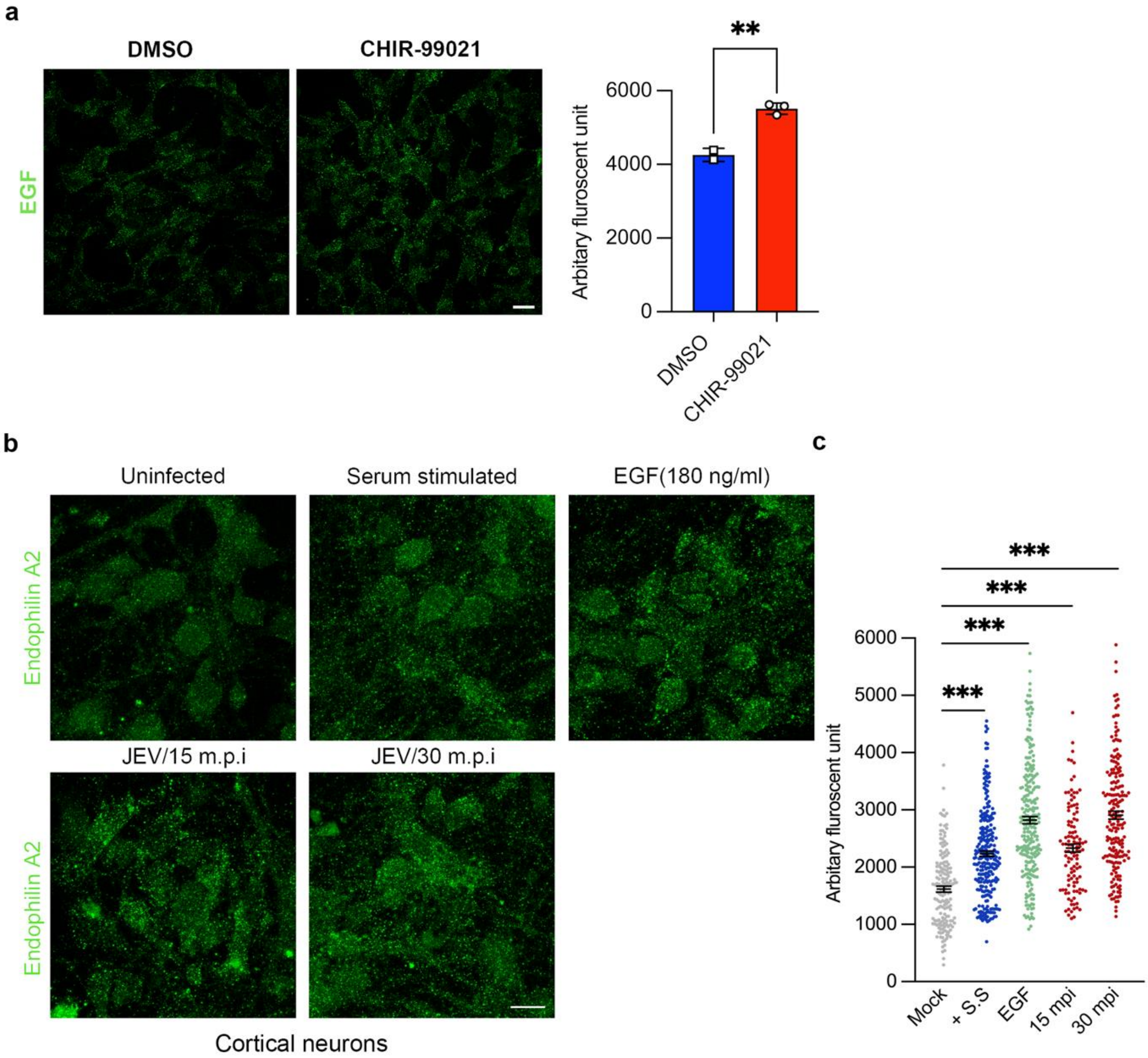


**Figure S4: Treatment with CHIR-99021 and serum stimulation upregulates FEME**. (a) SH-SY5Y cells were pre-treated with CHIR-99021 for 1 h at 37$^{O}$c and then pulsed with Alexa fluor 555 EGF (180 ng/ml) for 5 min at 37$^{O}$c. Cells were fixed, and images were acquired at a 63x objective Scale, 20 µm. Bar graph shows the quantification of the total fluorescent unit of EGF in DMSO and CHIR-99021 treated conditions. (b) Primary cortical neurons were left uninfected, 20% serum-stimulated, treated with EGF (180 ng/ml), or infected with JEV (100 MOI) for 15 min and 30 min respectively at 37$^{O}$c. Cells were fixed and stained with endophilin A2, and images were acquired using a 63x objective scale:10 µm. (c) The bar graph quantifies the total fluorescent unit of endophilin-positive puncta in different conditions. Analysis was performed using image J software with ~100 cells/coverslip. All values are represented as mean ± S.E.M; statistical analysis was determined with unpaired student's t-test/Kruskal-Wallis test with Dunn's multiple comparisons test. Statistical significance: NEJM: 0.12 (ns), 0.033 (*), 0.002(**), <0.001(***).

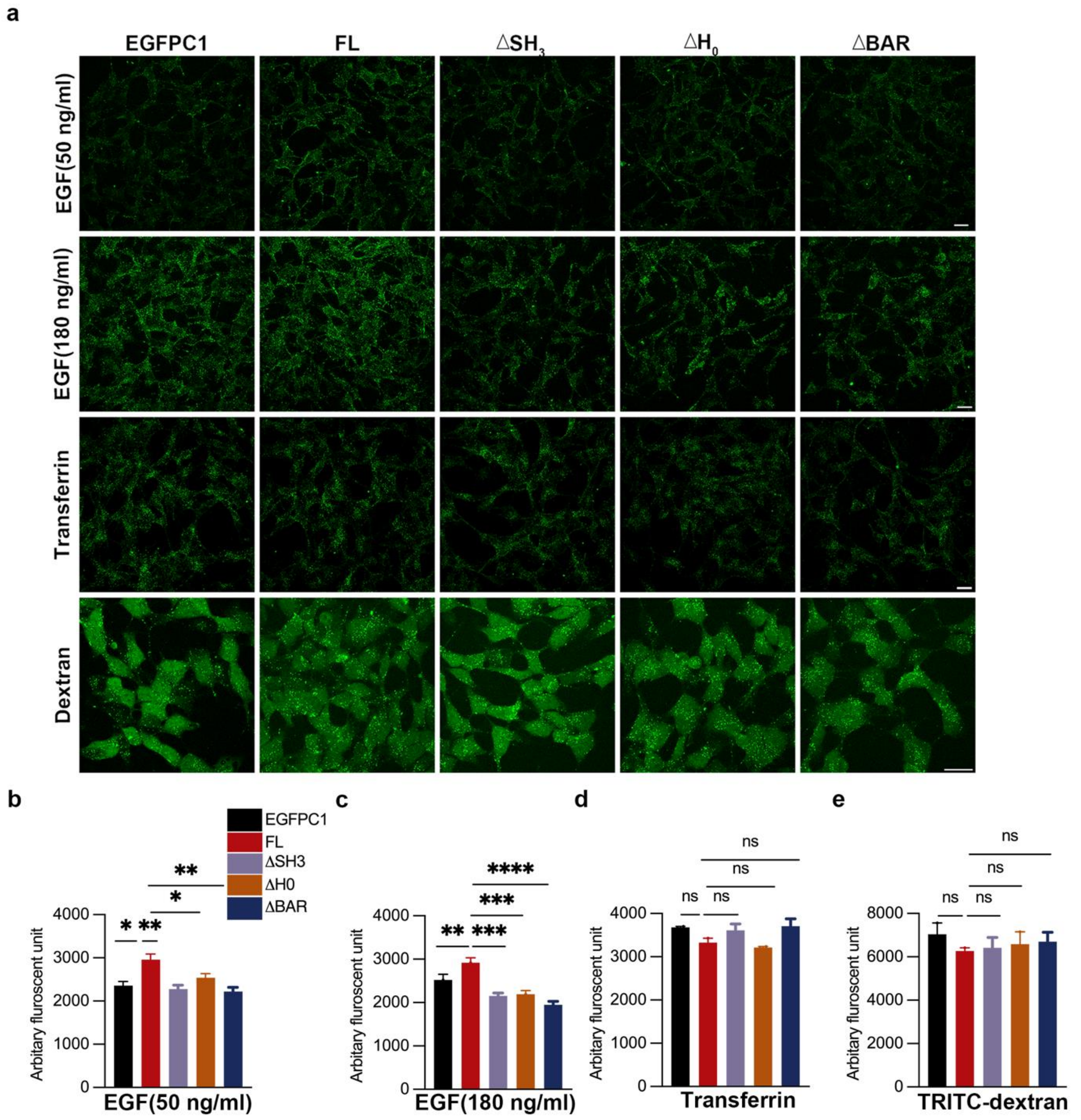


**Figure S5: Generation of SH-SY5Y clones expressing EGFP-tagged truncation domains of Endophilin.** (a) SH-SY5Y cells with stable expression of EGFPC1 (empty vector), endophilin A full length (FL), truncated endophilin with $\Delta SH_3$, $\Delta H_0$, and $\Delta$BAR domains were serum-stimulated (20% FBS), and pulsed with EGF-555 at 50 ng/ml or 180 ng/ ml for 5 min at $37^{O}$c. For the Tf uptake assay, cells were serum starved for 30 min and were pulsed with 20 ng/well of Tf-568 for 5 min at $37^{O}$c. Cells were pulsed with 200 ug/ml of TRITC- dextran (10,000 MW) for the fluid phase uptake assay for 10 min at $37^{O}$c. Cells were fixed with 4% PFA, and images were acquired with 63x objective. Images are representative of the uptake of fluorescently labelled cargoes. (b-e) The bar graph represents the quantification of total fluorescent intensities. Analysis was performed using image J software from ~100 cells per coverslip. All values are represented as mean ± S.E.M; statistical analysis was determined with Ordinary one-way ANOVA with Dunnett's multiple comparison test with 95% confidence level.

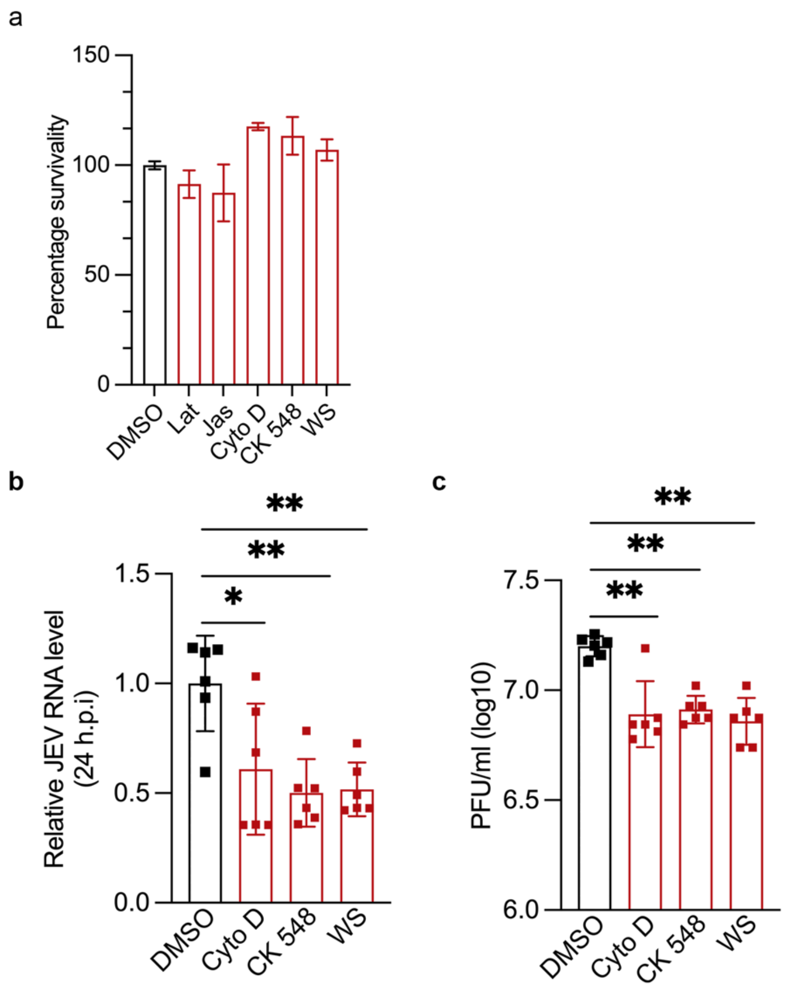


**Figure S6: Cell viability assay with pharmacological inhibitors of actin cytoskeleton.** (a) SH-SY5Y cells were treated with the actin inhibitors LatA (1uM), Jas (1uM), CytoD (5uM), CK-548 (50uM) and Wiskostatin (10uM) for 4 h, and a cell viability assay was performed to establish non-toxic drug concentration. (b-c) Cells were pre-treated with DMSO control/ actin inhibitor for 1 h at 37$^{O}$c, infected with 1 MOI virus at 37$^{O}$c for 1 h, and harvested at 24 hpi. Viral RNA was determined by qRT-PCR (b), and the extracellular virus particles were detected with plaque assays (c). Data shown are from two or more independent experiments represented as mean ± S.D. Statistical analysis was determined by Ordinary one-way ANOVA with Dunnett's multiple comparison test with 95% confidence level. Statistical significance: NEJM: 0.12 (ns),0.033 (*), 0.002(**), <0.001(***).

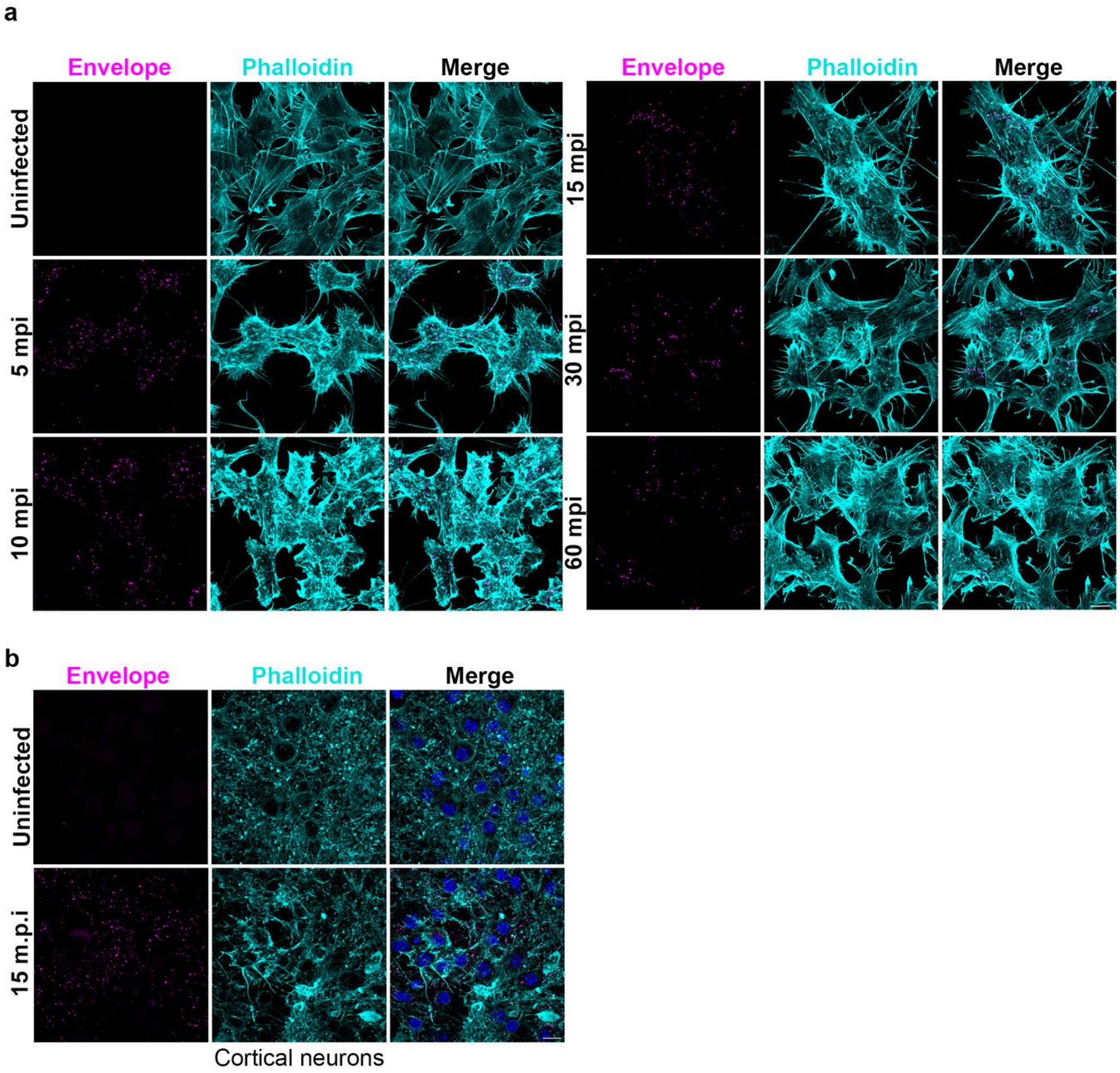


**Figure S7: Rearrangements in the F-actin organization upon JEV entry.** (a) SH-SY5Y cells were allowed to bind with 100 MOI JEV at $4^{O}$c to allow attachment only and were subsequently shifted to $37^{O}$c for internalization for 5-, 10-, 15-, 30-, 60- min. Cells were immunostained with JEV envelope antibody (magenta) and phalloidin (cyan) and imaged with Leica TCS SP8 microscope, 63x objective. The representative confocal micrograph shows the phalloidin-stained F-actin and JEV envelope puncta at the respective time points, scale: 10 µm. (b) Primary cortical neurons were infected with 100 MOI JEV at $4^{O}$c to allow attachment only and were subsequently shifted to $37^{O}$c for internalization for 15 min. Cells were fixed and immunostained with JEV envelope antibody (magenta) and phalloidin (cyan), nuclei with dapi (blue). The representative confocal micrograph shows the phalloidin-stained F-actin, envelope and nuclei in cortical neurons, scale: 10 µm.

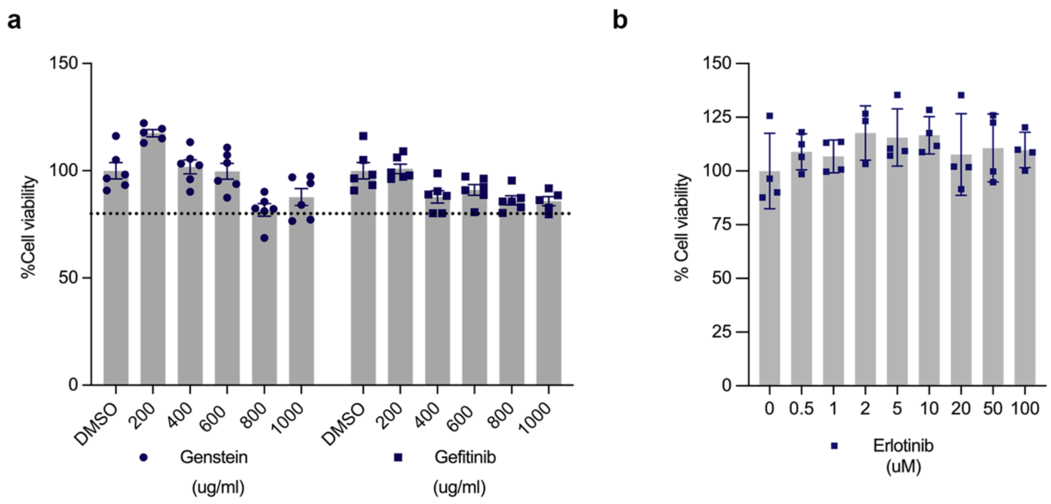


**Figure S8: Cell toxicity assay for genistein, gefitinib and erlotinib.** SH-SY5Y cells were either untreated or treated with genistein, gefitinib (a), or erlotinib (b) in a dose-dependent concentration for 5 h at 37$^{O}$c. MTT assay was performed to check the percentage of viable cells. Data shown are from two or more independent experiments represented as mean ± S.D.

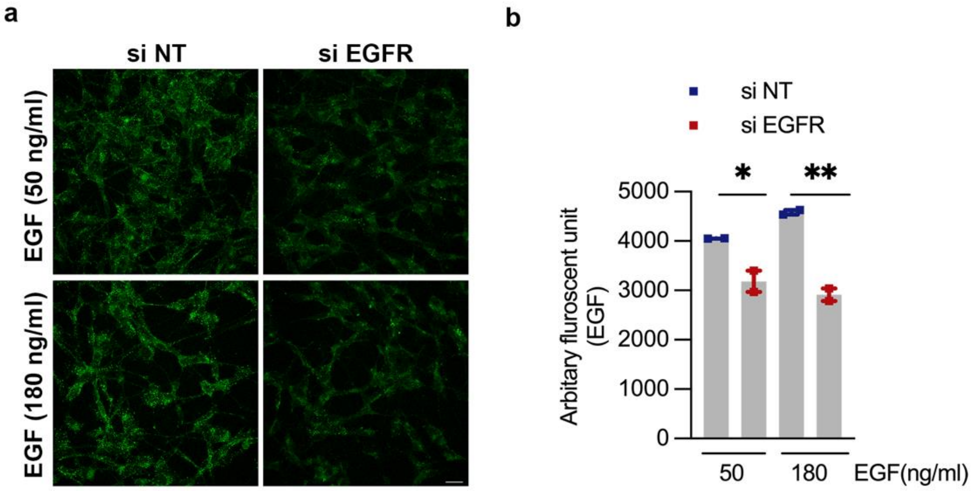


**Figure S9: EGFR silencing leads to a decrease in EGF uptake.** SH-SY5Y cells transfected with siNT/si EGFR for 72 h, were treated with Alexa fluor 555 EGF (50 & 180 ng/ml) for 5 min at $37^{O}$c. (a) Representative images indicating cargo uptake. Scale, 20 µm. (b) Bar graph shows the quantification of the total fluorescent units of EGF uptake. Analysis was performed using image J software with ~100 cells/coverslip, represented as mean ± S.E.M. Statistical analysis was determined with Ordinary one-way ANOVA with Šídák's multiple comparisons test, NEJM: 0.12 (ns),0.033 (*), 0.002(**), <0.001(***).

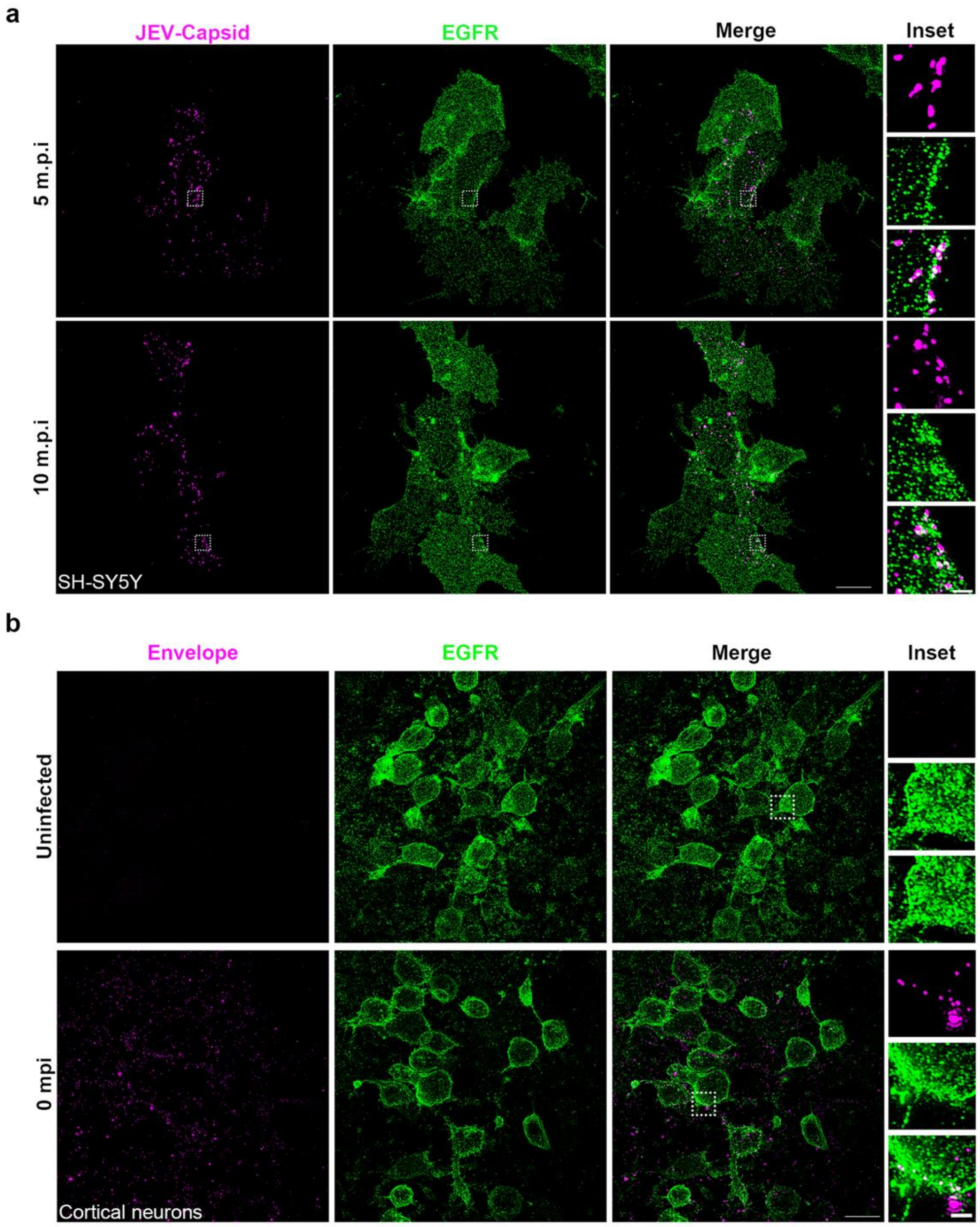


**Figure S10: EGFR colocalizes with the JEV capsid/envelope during entry.** (a) SH-SY5Y cells were allowed to bind with 100 MOI of virus for 1 h on ice and were fixed post 5- and 10-min internalization at 37$^{O}$c. Cells were immunostained with JEV capsid (magenta) and EGFR (green) antibodies. Images were acquired in Elyra PS1 (Carl Zeiss Super-resolution microscope) using 63 x objective. Insets show the zoomed image of capsid-positive structures colocalized with EGFR; scale: 10 µm, inset: 1 µm. (b) Primary cortical neurons were incubated with 100 MOI of virus for 1 h on ice and were subsequently fixed post 0-, 10-, 30-, and 60- min of virus internalization. Cells were immunostained with JEV envelope (magenta), and EGFR (green) antibodies. Images were acquired with Elyra PS1 (Carl Zeiss Super-resolution microscope) using 63 x objective; insets show the zoomed image of envelope-positive structures colocalized with EGFR. Representative image with 0 min pi is shown, data for other time points not included here; scale: 10 µm, inset: 1 µm.

**Table S1: Primer sequences used in the study**

| Genes | Forward (5'-3') sequence | Reverse (5'-3') sequence |
|---|---|---|
| hGAPDH | TGCACCACCAACTGCTTACG | GGCATGGACTGTGGTCATGAG |
| JEV | AGAGCACCAAGGGAATGAAATAGT TaqMan probe: CCACGCCACTCGACCCATAGACTG (5' end FAM, 3'end TAMRA). | AATAAGTTGTAGTTGGGCACTCTG |
| JEV Envelope region | TTACTCAGCGCAAGTAGGAGCGTCTCAAG | ATGCCGTGCTTGAGGGGGACG |
| SH3GL1 | GGGCAAGATCCCCGATGAG | CACCTGCTCGATGTCAGTCTC |
| SH3GL2 | CCAAACCTTCAGGTGTCCAAA | GCATCCCCTCATACCAGTTCTC |
| SH3GL3 | TTGGCTGTGTTCATAGAGGCA | TCGCATCTGTAGCTTGCTCTG |
| CLTB | CGAGGAGGCTTTCGTGAAGG | GCAGGCGGGACACATCTTT |
| EGFR | AGGCACGAGTAAGCTCAC | ATGAGGACATAACCAGCCACC |

**Table S2: Primer sequences used for cloning in the study**

| S. No. | Name | Sequence |
|---|---|---|
| 1. | FL-F | ATATAGATCTTCGGTGGCGGGGCTGAAG |
| 2. | FL-R | ATATGAATTCTCACTGCGGCAGGGGC |
| 3. | $\Delta H_O$-F | ATATAGATCTGGAGGGGCCGAGGGGAC |
| 4. | $\Delta H_O$-R | ATATGAATTCTCACTGCGGCAGGGGCA |
| 5. | $\Delta SH_3$-F | ATATAGATCTTCGGTGGCGGGGCTGAAG |
| 6. | $\Delta SH_3$-R | ATATGAATTCTCACTGGTCCAGGGGC |
| 7. | ΔBAR_1-F | ATATAGATCTATGTCGGTGGCGGGGC |
| 8. | ΔBAR_1-R | CGCTTAGGGCGTGAGGAGGCCCCTCCGA |
| 9. | ΔBAR_2-F | GTCGGAGGGGCCTCCTCACGCCCTAA |
| 10. | ΔBAR_2-R | ATATGAATTCTCACTGCGGCAGGGGCACA |

**Table S3: List of chemical reagents used the study**

| S. No. | Name of Reagent | Catalogue No. |
|---|---|---|
| 1. | Agarose Type VII | Sigma (A4018) |
| 2. | BCA assay kit | G-Biosciences (786-570) |
| 3. | Deoxyribonuclease I (DNase I) | SRL (61824) |
| 4. | Dimethyl sulfoxide (DMSO) | Sigma (276855-250ML) |
| 5. | EDTA | Sigma (E9884-500G) |
| 6. | Glucose | Sigma (G8270-100G) |
| 7. | HBSS | Gibco (14175095) |
| 8. | HEPES | Sigma (H3375-25G) |
| 9. | ImProm-II™ Reverse Transcription System | Promega (A3800) |
| 10. | Luminol | Santa Cruz sc-2048 |
| 11. | MTT 3-(4,5-Dimethylthiazol-2-yl)-2,5-Diphenyltetrazolium Bromide | VWR life science, (0793-1G) |
| 12. | Phenylmethylsulfonyl fluoride (PMSF) | Sigma (329-98-6) |
| 13. | Poly (ethylene glycol) (PEG 400) | Sigma (202398-500G) |
| 14. | Premix Ex Taq™ (Probe qPCR) | Takara (RR390A) |
| 15. | ProLong™ Gold Antifade Mountant with DAPI | Invitrogen (P36935) |
| 16. | Protease inhibitor cocktail (PI) | Sigma (P8340) |
| 17. | Puromycin | InvivoGen (ant-pr-1) |
| 18. | PVDF membrane | Merck Millipore (IPVH00010) |
| 19. | PhusionTM High-Fidelity DNA Polymerase | Thermo scientific (F530S) |
| 20. | QIAquick® PCR &Gel Cleanup kit | Qiagen (28506) |
| **S. No.** | **Name of Reagent** | **Catalogue No.** |
| 21. | Random hexamer | Sigma (H0268) |
| 22. | SDS | Sigma (L3771-500G) |
| 23. | Sodium pyruvate | HiMedia (TCL015) |
| 24. | SYBR® Premix Ex Taq™ | Takara (RR420A) |
| 25. | Triton™ X-100 | Sigma (T9284-500ML) |
| 26. | TRIzol reagent (RNAiso Plus) | Takara (9109) |
| 27. | Tween 20 | G-Biosciences (RC1227) |
| **Media & other additives** | | |
| 28. | 2XMEM | HiMedia (AL178A-500ML) |
| 29. | B-27 | Gibco (17504044) |
| 30. | DMEM | HiMedia (AL007A-500ML) |
| 31. | FBS | HiMedia, (RM10432-500ML) |
| 32. | Geneticin™ Selective Antibiotic (G418 Sulfate) | Gibco (10131035) |
| 33. | HiGlutaXL™ Dulbecco's Modified Eagle Medium | HiMedia (AL007G-500ML) |

| | | |
|---|---|---|
| 34. | L-15 | HiMedia (AL011S-500ML) |
| 35. | L-Glutamine | HiMedia (TCL012) |
| 36. | MEM | HyClone (SH3024401-500ML) |
| 37. | Neurobasal | Gibco (21103049) |
| 38. | Penicillin-Streptomycin | HiMedia (A007-100ML) |
| 39. | Trypsin - EDTA Solution | HiMedia (TCL007) |
| **Antibodies** | | |
| 40. | Chicken anti-Rabbit IgG (H+L) Cross-Adsorbed Secondary Antibody, Alexa Fluor 488, 1 mg | Invitrogen, A-21441 |
| 41. | Goat anti-Rabbit IgG (H+L) Highly Cross-Adsorbed Secondary Antibody, Alexa Fluor 647, 1 mg | Invitrogen, A-21245 |
| 42. | Donkey anti-Mouse IgG (H+L) Highly Cross-Adsorbed Secondary Antibody, Alexa Fluor 568, 1mg | Invitrogen, A10037 |
| 43. | Chicken anti-Mouse IgG (H+L) Cross-Adsorbed Secondary Antibody, Alexa Fluor 488, 1 mg | Invitrogen, A-21200 |
| 44. | Goat anti-Rabbit IgG (H+L) Cross-Adsorbed Secondary Antibody, Alexa Fluor™ 568 | Invitrogen, A-11011 |
| 45. | Goat anti-Mouse IgG (H+L) Cross-Adsorbed Secondary Antibody, Alexa Fluor™ 647 | Invitrogen, A-21235 |
| 46. | Alexa Fluor$^{TM}$ 555 EGF complex | Invitrogen (E35350), |
| 47. | Alexa Fluor$^{TM}$ 568 conjugate transferrin | Invitrogen (T23365) |
| 48. | Alexa Fluor$^{TM}$ 647 conjugate transferrin | Invitrogen (T23366) |
| 49. | Alexa Fluor™ 546 phalloidin | Invitrogen (A22283) |
| 50. | GAPDH | GeneTex (GTX100118) |
| 51. | JEV core protein C | GeneTex (GTX131368) |
| 52. | Endophilin I | Santa cruz (sc-134329) |
| 53. | Endophilin II | Santa cruz (sc-365704) |
| 54. | Endophilin III | Santa cruz(sc-376592) |
| 55. | Clathrin light chain | Abcam (ab129326), |
| 56. | EGFR | Santa cruz (sc-120) |
| 57. | EGFR | CST, 2232 |
| 58. | Phospho-EGF Receptor (Tyr1068) | CST, 2234 |
| 59. | Akt | CST, 9272 |
| 60. | Phospho-Akt (Ser473) | CST, 9271 |
| 61. | Peroxidase AffiniPure Donkey Anti-Mouse IgG (H+L) | Jackson ImmunoResearch (715-035-150) |
| 62. | Peroxidase AffiniPure Donkey Anti-Rabbit IgG (H+L) | Jackson ImmunoResearch (711-035-152) |

| | | |
|---|---|---|
| 63. | GFP | CST (2555) |
| 64. | JEV-Envelope | Abcam (ab41671) |
| **Proteins** | | |
| 65. | Human EGF | Sigma (E9644) |
| 66. | Recombinant Human EGFR Protein (ECD, hFc Tag) | Sino Biological, 10001-H02H |
| 67. | Recombinant Human EGFR Protein (Isoform Viii, hFc Tag) | Sino Biological, 29662-H02B |
| **siRNA/Transfection reagents** | | |
| 68. | DharmaFECT 1 | T-2001-03 |
| 69. | Lipofectamine$^{TM}$ RNAimax | Invitrogen (13778030) |
| 70. | Lipofectamine™ 2000 | Invitrogen (11668019) |
| 71. | ON-TARGET plus Non-targeting (NT) | Dharmacon (D-001810-10-20) |
| 72. | ON-TARGET plus human SH3GL1 | Dharmacon (L-019582-00-0005) |
| 73. | ON-TARGET plus human SH3GL2 | Dharmacon (L-012597-00-0005) |
| 74. | ON-TARGET plus human SH3GL3 | Dharmacon (L-015728-02-0005) |
| 75. | ON-TARGET plus human CLTB | Dharmacon (L-004003-00-0010) |
| 76. | ON-TARGET plus human EGFR | Dharmacon (L-003114-00-0005) |
| **Small inhibitors** | | |
| 77. | Erlotinib | HY-50896 |
| 78. | Genistein | 92136-10MG |
| 79. | Gefitinib | SML1657-10MG |
| 80. | Cetuximab | Selleckchem, A2000 |
| 81. | Dynasore | D7693 |
| 82. | CHIR-99021 | 4423 |
| 83. | Cytochalasin D (CytoD) | Sigma (CytoD, C8273) |
| 84. | Latrunculin A (Lat A) | Sigma (Lat A, L5163) |
| 85. | Jasplankinolide (Jas) | Sigma (Jas, J4530) |
| 86. | CK 548 | Sigma (C7499) |
